# SUMO-specific protease 2 (SENP2) suppresses browning of white adipose tissue through C/EBPβ modulation

**DOI:** 10.1101/2020.12.16.422969

**Authors:** Ji Seon Lee, Sehyun Chae, Jinyan Nan, Young Do Koo, Seung-Ah Lee, Young Joo Park, Daehee Hwang, Weiping Han, Young-Bum Kim, Sung Soo Chung, Kyong Soo Park

**Affiliations:** Department of Internal Medicine, Seoul National University College of Medicine, Seoul, Korea; Korea Brain Bank, Korea Brain Research Institute, Daegu, Korea, Daegu, Korea; Department of Molecular Medicine and Biopharmaceutical Sciences, Graduate School of Convergence Science and Technology, College of Medicine, Seoul National University, Seoul, Korea; Metabolism in Human Diseases Unit, Institute of Molecular and Cell Biology, Agency for Science, Technology and Research (A*STAR), Singapore, Singapore; School of Biological Sciences, Seoul National University, Seoul, Korea; Division of Endocrinology, Diabetes and Metabolism, Department of Medicine, Beth Israel Deaconess Medical Center and Harvard Medical School, Boston, MA, USA; Biomedical Research Institute, Seoul National University Hospital, Seoul, Korea

**Author notes:** Corresponding authors: (Lead Contact) Kyong Soo Park, Department of Internal Medicine, Seoul National University College of Medicine, Seoul, Korea, 03080, Sung Soo Chung, Biomedical Research Institute, Seoul National University Hospital, Seoul, Korea, 03082. Summary* - Adipocyte-specific *Senp2* knockout (*Senp2*-aKO) promotes WAT browning. - *Senp2*-aKO mice are protected from high fat-induced obesity and insulin resistance. - Reduction of *Hoxc10* transcription is essential for SENP2–depletion-mediated beige adipogenesis in iWAT. - Sumoylated C/EBPβ efficiently suppresses *Hoxc10* transcription by recruiting the corepressor DAXX. - C/EBPβ mediates thermogenic adipocyte differentiation by SENP2 depletion in both iWAT and eWAT.

## Abstract

SUMO-specific protease 2 (SENP2) is highly expressed in white adipose tissue (WAT) and plays an important role in the early stages of adipogenesis. To investigate the function of SENP2 in adipocytes, we generated adipocyte-specific *Senp2* knock-out (*Senp2*-aKO) mice. Compared to wild-type mice, *Senp2*-aKO mice had reduced adipose tissue mass and smaller multi-locular adipocytes in inguinal WAT (iWAT). Body temperatures of *Senp2*-aKO mice were effectively regulated during cold exposure. Additionally, *Senp2*-aKO mice were resistant to high–fat–diet-induced obesity and insulin resistance and exhibited an increase in energy expenditure rates. Expression of thermogenic genes, including *Ucp1*, was significantly increased in iWAT (and less efficiently in epidydimal WAT [eWAT]) of *Senp2*-aKO mice, suggesting that SENP2 depletion accelerates browning of WAT. Further, suppression of HOXC10 was essential for beige adipocyte formation in SENP2-deficient cells of iWAT, and *Hoxc10* transcriptional suppression was mediated by C/EBPβ, a direct target of SENP2. Sumoylated C/EBPβ efficiently inhibited *Hoxc10* transcription through recruitment of the transcriptional co-repressor DAXX. Similarly, *Senp2* knockdown using siRNAs during adipogenesis promoted thermogenic adipocyte differentiation of precursor cells in both iWAT and eWAT, and C/EBPβ was a common mediator. Together these results suggest that SENP2 plays critical role in white adipocyte differentiation by suppressing differentiation toward thermogenic adipocytes through modulation of C/EBPβ in both iWAT and eWAT.

## Introduction

Obesity is the primary contributing factor in several metabolic diseases and is caused by expansion of white adipose tissue (WAT) (1). White adipocytes consist of a single, large lipid droplet that stores energy, while brown adipocytes are responsible for adaptive thermogenesis (2, 3). Uncoupling protein 1 (UCP1), which is specifically expressed in brown adipocytes, catalyzes a proton leak across the mitochondrial membrane that results in heat generation (4). Recently, a third class of adipocytes, beige adipocytes, was identified as clusters of cells embedded in WAT. Beige adipocytes have multi-locular lipid droplets, similar to those of brown adipocytes, and contribute to thermogenesis through expression of UCP1 (5–7). Therefore, the induction of beige adipocytes is a promising therapeutic approach for obesity and diabetes.

Beige adipocyte formation in WAT is activated by cold temperature, β-adrenergic stimulation, chronic exercise, and various pharmacological treatments (8–13), and can also drive from mature white adipocytes (14). Conversely, beige adipocytes can switch to white adipocytes after an initial cold exposure ends but retain genetic memory and convert back to beige adipocytes upon a second cold stimulation (15). Another source of beige adipocytes is WAT progenitors. Several studies indicate that progenitor cells expressing specific markers, such as platelet-derived growth factor receptor beta, alpha smooth muscle actin and myosin heavy chain 11, generate beige adipocytes upon cold exposure (16–20). However, further studies are needed to understand the relative contributions of these cells to beige adipocyte formation.

Several transcription factors are involved in the adipose browning process (21, 22). First, peroxisome proliferator-activated receptor γ (PPARγ) coactivator 1α (PGC1α) is induced by cold exposure and activates expression of thermogenic genes (23). Loss of PGC1α in WAT reduces UCP1 expression and thermogenesis (24). PRDM16, a member of the PR/SET domain family, is highly expressed in brown adipose tissue (BAT), and controls brown fat development by binding and enhancing the transcriptional activities of CCAAT/enhancer binding protein β (C/EBPβ), PPARγ and C2H2 zinc-finger protein 516 (22, 25). Subcutaneous WAT (scWAT) is very susceptible to browning, and PRDM16 is highly expressed in subcutaneous fat compared to visceral fat depots. Adipocyte-specific depletion of PRDM16 induces defects in scWAT thermogenesis and the conversion of subcutaneous to visceral fat (26). Further, overexpression of PRDM16 increases beige adipocytes in scWAT and induces protective effects against high–fat–diet (HFD)-induced obesity and insulin resistance (27).

Genes of the homeobox (*Hox*) family exhibit specific expression patterns in both WAT and BAT (28, 29). For example, *Hoxc8*, *Hoxc9* and *Hoxc10* are highly expressed in WAT, while differentiation of human WAT progenitor cells into beige adipocytes is associated with downregulation of *Hoxc8* (30). Expression of both *Hoxc9* and *Hoxc10* is higher in subcutaneous than visceral adipose tissue and their expression degree is correlated with obesity (31). Furthermore, ectopic expression of *Hoxc10* suppresses cold-induced browning in scWAT, suggesting that HOXC10 is a key negative regulator of browning (32).

In our previous studies, we showed that small ubiquitin-like modifier (SUMO)-specific protease 2 (SENP2) is required for adipogenesis and is involved in lipid metabolism in skeletal muscle. Specifically, knockdown of *Senp2* expression in 3T3L1 preadipocytes inhibited adipogenesis through regulation of C/EBPβ stability during the early stages of adipogenesis (33). *Senp2* expression was positively regulated by palmitate, and overexpression of *Senp2* in skeletal muscle protected mice from HFD-induced obesity and insulin resistance by increasing fatty acid oxidation (FAO). SENP2 enhanced the transcription of genes encoding FAO-associated enzymes, including long-chain acyl-CoA synthetase 1 and carnitine-palmitoyl transferase 1b, through desumoylation-mediated activation of PPARδ and PPARγ (34).

In the current study, we generated adipocyte-specific *Senp2* knock out mice (*Senp2*-aKO) to thoroughly investigate the physiological roles of SENP2 in adipocytes. Interestingly, knockout of *Senp2* using an adiponectin-Cre system enhanced browning of iWAT in the absence of cold exposure or treatment with a browning inducer. Additionally, *Hoxc10* transcription was significantly reduced in the iWAT of *Senp2*-aKO mice, allowing the differentiation of white adipose progenitor cells toward thermogenic adipocytes. Furthermore, accumulation of sumoylated C/EBPβ, a direct target of SENP2, efficiently suppressed *Hoxc10* transcription. These results suggest that SENP2 negatively regulates browning and is important for maintaining white adipocyte identity during the differentiation of white adipose progenitor cells.

## Results

### Generation of adipocyte-specific Senp2 knockout mice (Senp2-aKO)

SENP2 expression was relatively high in white adipose tissues, inguinal WAT (iWAT) and epididymal WAT (eWAT), compared to BAT. Much lower levels of SENP2 were detected in muscle and liver, suggesting an important role of SENP2 in adipose tissues (Supplemental Fig. 1A). To investigate the function of SENP2 in adipocytes, we generated an adipocyte-specific *Senp2* knockout (*Senp2*-aKO) mouse model in which exon 3 of *Senp2* was deleted by Cre recombinase driven by the adiponectin promoter (Fig. 1A). SENP2 protein and mRNA levels were dramatically reduced in iWAT, eWAT, and BAT in *Senp2*-aKO (aKO) mice compared to control mice (Fig. 1B and Supplemental Fig. 1B). When adipocytes were separated from the stromal vascular fractions (SVF) in iWAT, SENP2 protein levels were significantly reduced in adipocytes of *Senp2-aKO* mice compared with control mice, but were not changed in the SVF (Supplemental Fig. 1C). In addition, SENP2 expression in muscle or liver tissue was not changed in *Senp2*-aKO mice compared with control mice (Supplemental Fig. 1D).

**Figure 1.**
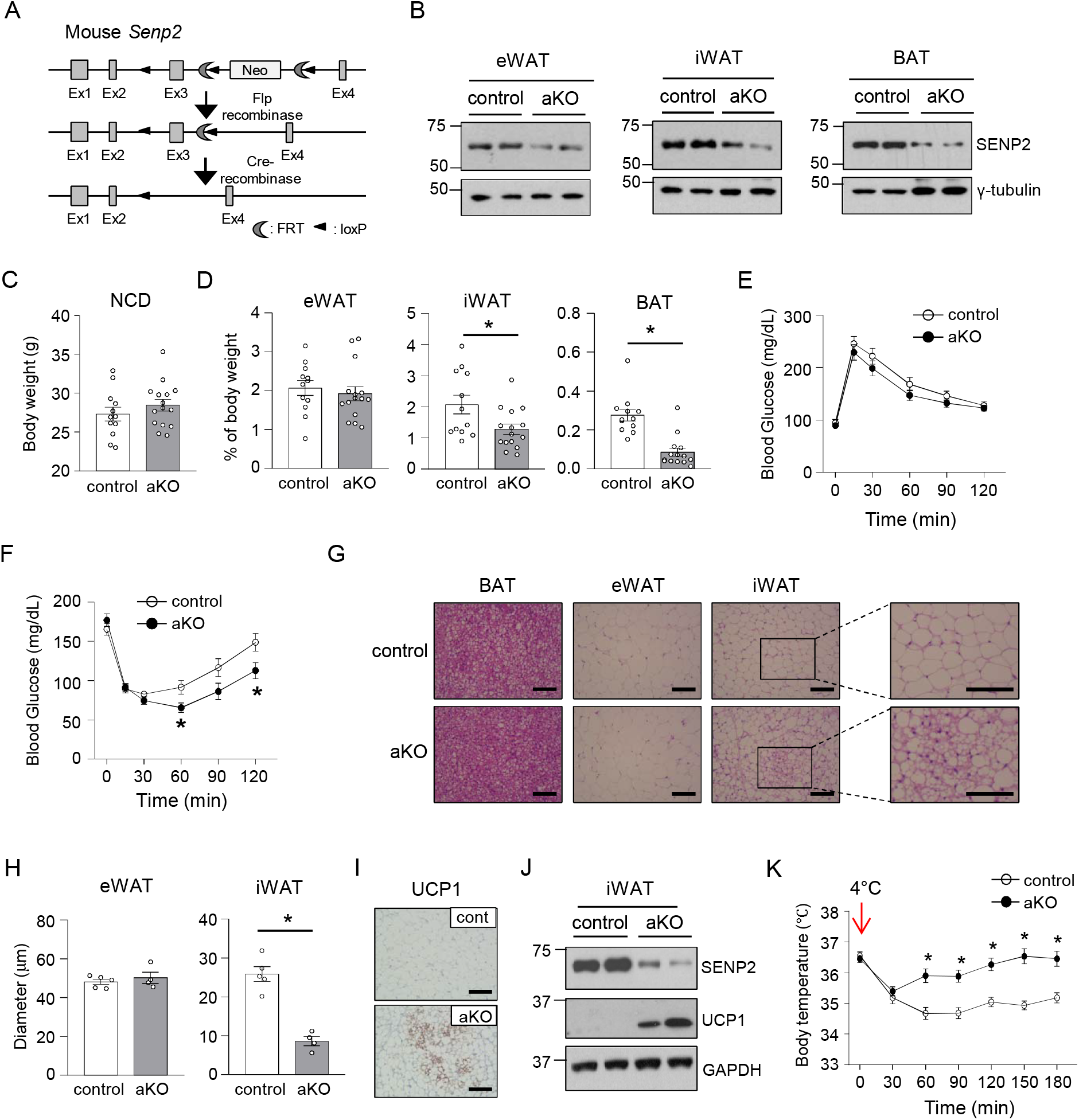
Accumulation of beige adipocytes in iWAT of *Senp2*-aKO mice. (A) Generation of *Senp2*-aKO mice. (B) Expression of SENP2 in adipose tissues of *Senp2*-floxed mice (control) and *Senp2*-aKO mice (aKO). (C-L) Control and *Senp2*-aKO mice were fed a normal chow diet (NCD) for 16 weeks. (C) Body weights of 16 week-old control and *Senp2*-aKO mice. n = 12-15. (D) Each fat pad weight was expressed as % of each body weight. \**P* < 0.05 by Student’s t-test, n = 11-15. (E) Glucose tolerance test (GTT) and (F) insulin tolerance test (ITT) \**P* < 0.05 by Student’s t-test, n = 10-12. (G) Representative H&E staining of eWAT, iWAT, and BAT from control and *Senp2*-aKO mice. Scale bar, 100 μm. (H) Average size of adipocytes in eWAT and iWAT. \**P* < 0.05 by Student’s t-test, n = 4-5. (I) Immuno-histochemical staining of iWAT using an UCP1 antibody. Scale bar, 100 μm. (J) Western blot analysis using an UCP1 antibody. (K) Control and *Senp2*-aKO mice (16-17-week old) were exposed to cold temperature (4°C) for 3 h. Body temperature was measured every 30 min. \**P* < 0.05 by Student’s t-test, n = 10-12. All data are presented as mean ± SEM.

### Adipocyte-specific depletion of SENP2 improved thermogenesis through accumulation of beige adipocytes in iWAT

When mice were fed a normal chow diet, we observed no difference in body weights between control and *Senp2*-aKO mice (Fig. 1C). However, both iWAT and BAT mass significantly decreased in *Senp2*-aKO mice compared to control mice, whereas eWAT mass did not change (Fig. 1D). We observed no significant difference in glucose tolerance between control and *Senp2*-aKO mice, but *Senp2*-aKO mice had an improved insulin response (Fig. 1E and 1F). Histological analysis of adipose tissues from 12-week old mice showed that the average size of iWAT adipocytes from *Senp2*-aKO mice was smaller than that of control mice, whereas we found no change in eWAT adipocytes between the two groups (Fig. 1G and 1H). In addition, iWAT from *Senp2*-aKO mice had abundant multi-locular adipocytes and UCP1 expression (Fig. 1G and 1I-J). To compare adaptive thermogenesis, control and *Senp2*-aKO mice were exposed to cold conditions (4°C). Body temperatures of both groups of mice dramatically decreased 30 min after cold exposure. However, unlike control mice, *Senp2*-aKO mice gradually recovered their normal body temperature (Fig. 1K). Taken together, these results suggest that beige adipocytes accumulate in iWAT of *Senp2-*aKO mice and contribute to maintaining normal body temperature in a cold environment.

### Adipocyte-specific depletion of SENP2 protects mice from HFD-induced obesity and insulin resistance

When mice were fed a HFD, body weight gain was attenuated in *Senp2*-aKO mice compared to control mice although there was no change in food intake by the mice (Fig. 2A and Supplemental Fig. 2A). The total body fat mass of the *Senp2*-aKO mice was significantly lower (~40% reduction) than that of the control mice, but no change was observed in the lean mass or bone mineral density (Fig. 2B and Supplemental Fig. 2B). The average weight of each adipose tissue (eWAT, iWAT, and BAT) in *Senp2*-aKO mice was significantly lower than those in control mice, whereas we saw no change in the weight of other tissues, such as liver and muscle (Fig. 2C and Supplemental Fig. 2C).

**Figure 2.**
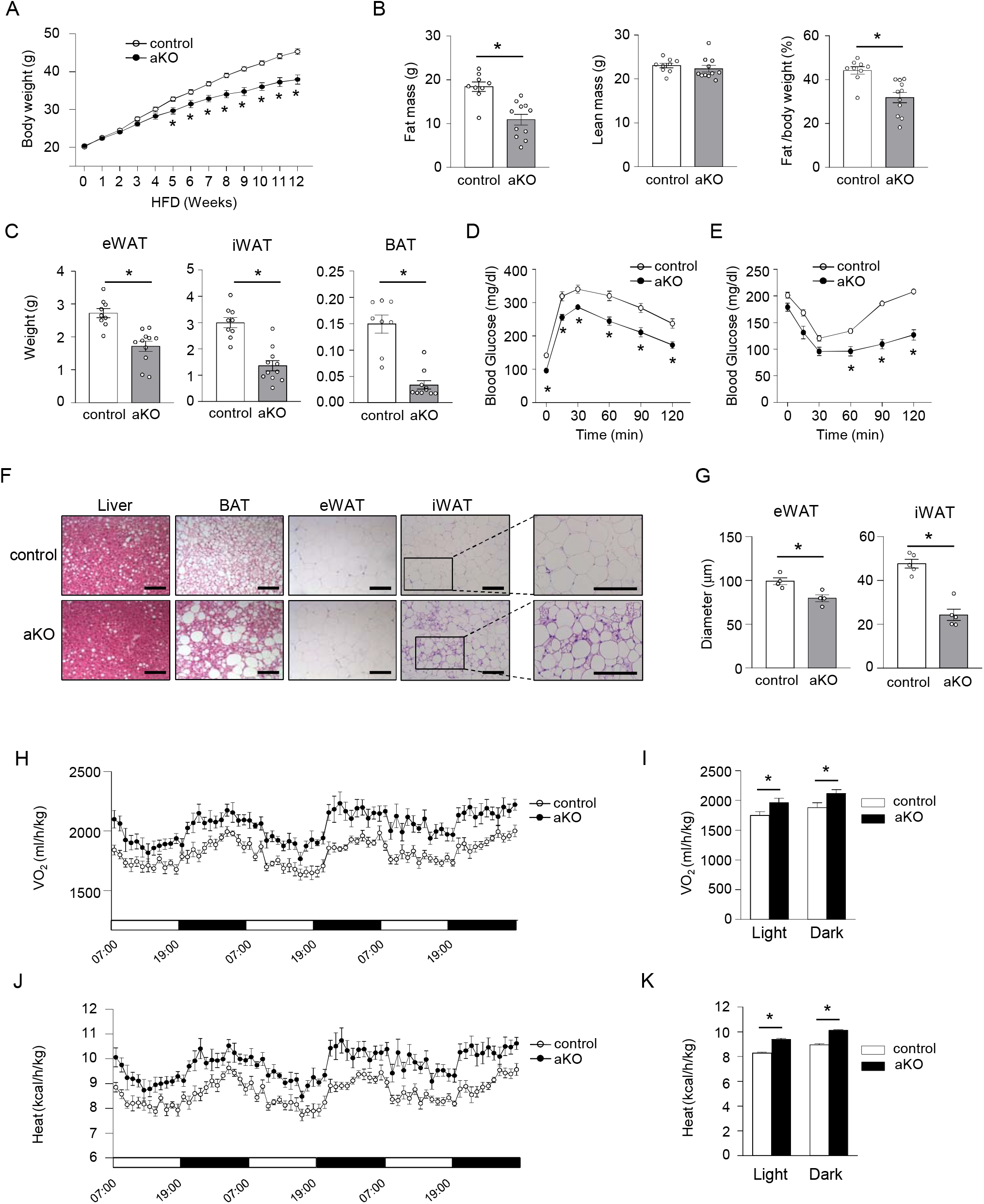
Adipocyte-specific depletion of SENP2 ameliorates HFD-induced obesity and insulin resistance. (A) Body weights of control and *Senp2*-aKO mice were measured during 12 weeks of HFD feeding. \**P* < 0.05 by Student’s t-test, n =19. (B) Fat mass and lean mass were measured using DEXA after 14 weeks of HFD feeding (left and middle panels), and % fat weight/body weight was also shown (right panel). \**P* < 0.05 by Student’s t-test, n = 9-11 (C) Fat pad weight of control and *Senp2*-aKO mice was measured after 14 weeks of HFD feeding. \**P* < 0.05 by Student’s t-test, n = 8-11. (D) GTT and (E) ITT after 12 or 13 weeks of HFD feeding, respectively. \**P* < 0.05 by Student’s t-test, n = 13-15. (F) Representative H&E staining of WATs, BAT and liver tissues after 14 weeks of HFD feeding. The right panel shows enlarged photos of iWAT. Scale bar, 100 μm. (G) Average size of adipocytes in eWAT and iWAT. \**P* < 0.05 by Student’s t-test, n = 4-5. (H) Volume oxygen (O_2_) consumption rates and (J) heat production rates at week 15 of HFD-feeding. (I and K) Average O_2_ consumption and average heat production of 3 day measurements. \**P* < 0.05 by Student’s t-test, n = 10-12. All data are presented as mean ± SEM.

To determine whether adipocyte-specific SENP2 depletion affected HFD-induced insulin resistance, glucose tolerance tests and insulin tolerance tests were performed after 12 or 13 weeks of HFD feeding. *Senp2*-aKO mice showed improved glucose tolerance, as well as decreased blood glucose levels in response to insulin (Fig. 2D and 2E). Serum insulin, triglyceride and free fatty acid levels were similar in control and *Senp2*-aKO mice (Supplemental Fig. 2D and 2E). In the *Senp2*-aKO mice, serum leptin and adiponectin levels were lower than those of control mice (Supplemental Fig. 2F), which was probably due to reduced fat mass. We also investigated morphological differences in adipose tissue between HFD-fed control and *Senp2*-aKO mice. Adipocyte size in WAT (eWAT and iWAT) was significantly reduced in *Senp2*-aKO mice (Fig. 2F and 2G). Further, we easily observed multi-locular droplets in iWAT of *Senp2*-aKO mice (Fig. 2F, right panel). In contrast, large lipid droplets were detected in BAT of *Senp2*-aKO (Fig. 2F).

Next, we measured energy expenditure rates to investigate the mechanism underlying the effects of *Senp2* knockout and found that O_2_ consumption and heat production rates increased in *Senp2*-aKO mice during both light and dark cycles (Fig. 2H–2K). These results suggest that adipocyte-specific SENP2 depletion elevates energy expenditure, which results in protection of mice from HFD-induced obesity and insulin resistance.

### SENP2 depletion promotes expression of thermogenic genes in iWAT and eWAT

The iWAT gene expression profiles of HFD-fed control and *Senp2*-aKO mice were analyzed using micro-arrays and quantitative PCR (qPCR) analysis. Genes involved in negative regulation of lipid storage, heat generation and brown fat differentiation were up-regulated in iWAT of *Senp2*-aKO mice, whereas genes related to steroid or lipid biosynthesis were down-regulated in *Senp2*-aKO iWAT (Fig. 3A). While the mRNA levels of genes abundantly expressed in white adipocytes were reduced or unchanged, expression of brown–adipocyte–specific genes, such as *Ucp1, Cidea, Elovl3,* and *Dio2*, was significantly elevated in iWAT of *Senp2*-aKO mice (Fig. 3B and 3C). The difference in expression of BAT-specific genes between HFD-fed control and *Senp2*-aKO mice was greater than the difference seen in mice fed a normal chow diet (Fig. 3C and 3D; 30-fold vs. 4-fold for *Ucp*1). The mRNA levels of mitochondrial oxidative–phosphorylation–related genes, such as *Cox7a1* and *Cox8b*, were also higher in iWAT of *Senp2*-aKO mice (Fig. 3C and 3D). These results indicate the efficient accumulation of beige adipocytes in iWAT by SENP2 deficiency.

**Figure 3.**
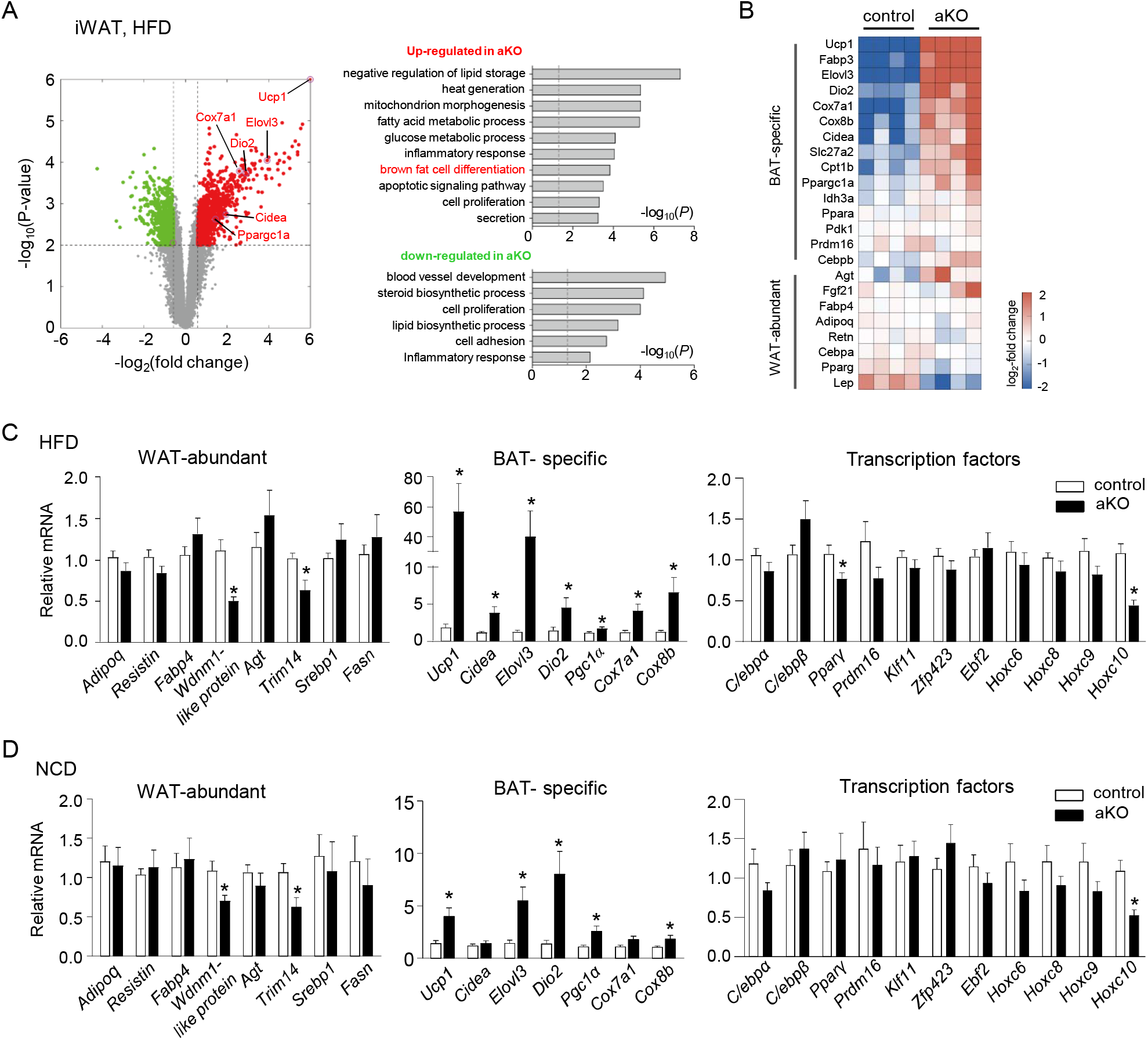
SENP2 depletion induced thermogenic gene expression in iWAT. (A) Volcano plot (left) showing up- (red) and down-regulated (green) genes in iWAT from *Senp2*-aKO mice, compared to that from control mice on a HFD for 14 weeks. Cellular processes enriched by the up-(top) and down-regulated (bottom) genes are also shown in the bar plots (right) where the enrichment significance (p-value) was displayed as -log_10_(p). (B) Heat map showing the log_2_-fold changes of the indicated genes with respect to their median mRNA expression levels in control and *Senp2*-aKO iWAT. n = 4. (C) Relative gene expression in iWAT of HFD-fed mice was determined by qPCR. The mRNA level of each gene in the control mice were set to 1. \**P* < 0.05 by Student’s t-test, n = 14-15. (D) Relative gene expression in iWAT of NCD-fed mice. \**P* < 0.05 by Student’s t-test, n = 13-14. All data are presented as mean ± SEM.

We compared the expression levels of several transcription factors related to adipogenesis and browning and saw no significant change of those tested, including *Prdm16*, an important regulator that induces browning (Fig. 3C and 3D, right panel and Supplemental Fig. 3A). However, expression of *Hoxc10*, a known negative regulator of browning, was significantly reduced in iWAT of HFD-fed *Senp2*-aKO mice (Fig. 3C) which was consistently observed under normal chow-diet feeding (Fig. 3D), suggesting the possibility that reduced expression of *Hoxc10* accelerates browning of iWAT in *Senp2*-aKO mice.

We also compared gene expression patterns of eWAT in control and *Senp2*-aKO mice. Expression of *Ucp1*, *Elovl3* and *Dio2* in eWAT of *Senp2*-aKO mice was higher than that of control mice fed either a normal chow diet or HFD (9–15 fold for *Ucp1*, 2–5 fold for *Elovl3* and *Dio2*) (Supplemental Fig. 3B and 3C). Unlike iWAT, *Hoxc10* mRNA levels in eWAT were very low (Supplemental Fig. 3D) and were not affected by *Senp2* knockout (Supplemental Fig. 3B and 3C). Although beige-like adipocyte depots were not observed in eWAT based on the H&E staining results (Fig. 2F), the increased thermogenic gene expression seen in eWAT may partially contribute to alleviating HFD-induced obesity and insulin resistance in *Senp2*-aKO mice.

### SENP2 depletion facilitates differentiation of white adipose progenitor cells into thermogenic adipocytes

We found that *Senp2* knockout results in thermogenic gene expression in white adipocytes, therefore we next accessed to figure out the mechanism to induce thermogenic adipocytes. The SVF was isolated from iWAT of control and *Senp2*-aKO mice, and then differentiation into adipocytes was induced (Fig. 4A). After differentiation, lipid droplets were stained with Oil-Red O (Fig. 4B). Similar to the gene expression patterns in iWAT, expression of brown– adipocyte–specific genes was higher (5–6 fold for *Ucp1* and *Cidea*; 2–3 fold for *Elovl3*, *Dio2* and *Pgc1*α), while *Hoxc10* expression was lower in SVF-derived adipocytes from *Senp2*-aKO mice (Fig. 4C). To confirm the thermogenic response of the SENP2-deficient adipocytes, cells were treated with the β3-adrenergic receptor agonist CL316243. *Ucp1* and *Pgc1*α mRNA levels were rapidly increased upon CL316243 treatment in SENP2-deficient adipocytes but not in control adipocytes (Fig. 4D). Increased UCP1 expression was confirmed by western blot analysis (Fig. 4E). We also compared gene expression patterns at several time points during differentiation of the progenitor cells into adipocytes. In the cells isolated from *Senp2*-aKO mice, *Senp2* mRNA levels were decreased at day 4, likely responding to increased adiponectin. *Hoxc10* mRNA levels also decreased, whereas *Ucp1* mRNA levels gradually increased in SENP2-deficeint cells (Fig. 4F). Similar to *Ucp1*, expression of *Cidea*, *Elovl3* and *Pgc1α* also gradually increased in SENP2-deficient cells during adipogenesis (Supplemental Fig. 4). These results suggest that SENP2 depletion during adipogenesis of white adipose progenitor cells changes the direction of differentiation toward beige or beige-like adipocytes.

**Figure 4.**
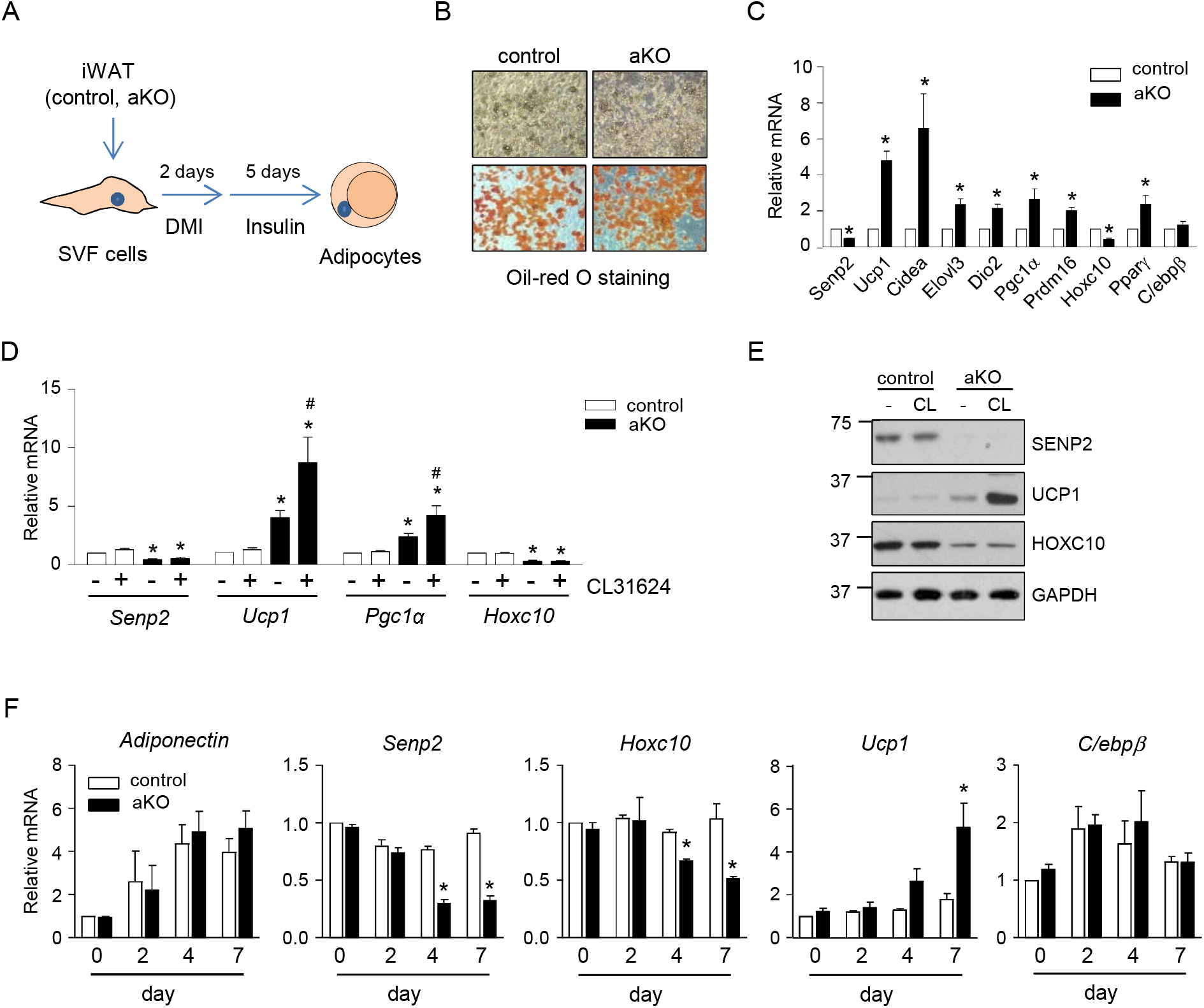
SENP2 depletion facilitates differentiation of white adipose progenitor cells toward thermogenic adipocytes. (A-C) SVF was isolated from iWAT of control and *Senp2*-aKO mice and then differentiated to adipocytes. (B) Oil-Red O staining at day 7 of differentiation. (C) Relative mRNA levels of BAT specific genes and some transcription factors at day 7 of differentiation. The values obtained in the cells from control mice were set to1. \**P* < 0.05 by Student’s t-test, n = 9-10. (D-E) At day 7 of differentiation, CL31624 (10 μmol/L) were treated for 5 h. The mRNA levels were measured (D), and the values in the cells from control mice without CL31624 treatment were set to 1 and the others were its relative values. \**P* < 0.05 vs control/without CL31624 by Student’s t-test, ^#^ *P* < 0.05 vs aKO/without CL31624 by 2-way Anova. n = 4. (E) Western blotting was performed. (F) At different points of time (0, 2, 4 and 7 day) after the induction of adipocyte differentiation from SVF, RNAs were isolated and subjected to qPCR. The values of control cells at day 0 were set to 1 and the others were its relative values. \**P* < 0.05 vs control by 2-way Anova, n = 3. All data are represented as the mean ± SEM.

### Suppression of Hoxc10 is necessary for Senp2 knockout-induced beige adipogenesis in iWAT

HOXC10 is a known negative regulator of browning (32), and *Hoxc10* expression was suppressed in iWAT of *Senp2*-aKO mice and their adipocytes (Fig. 3C-D and 4C). Therefore, we tested whether *Hoxc10* overexpression inhibited *Senp2*–knockout-induced beige adipocyte differentiation. Five days after the induction of adipocyte differentiation in iWAT SVF cells of control or *Senp2*-aKO mice, cells were infected with adeno-associated virus expressing *Hoxc10* (AAV-*Hoxc10*) (Fig. 5A). We confirmed *Hoxc10* overexpression at both the mRNA and protein levels (Fig. 5B and Supplemental Fig. 5A). UCP1 induction in SENP2-deficient adipocytes was completely inhibited by overexpression of HOXC10 (Fig. 5B). In addition to *Ucp1*, expression of other brown–adipocyte–specific genes, including *Cidea*, *Elovl3*, *Dio2*, and *Pgc1a*, were not induced in SENP2-deficient adipocytes after *Hoxc10* overexpression (Fig. 5C), indicating that suppression of *Hoxc10* expression is necessary for beige adipocyte differentiation in iWAT of *Senp2*-aKO mice.

**Figure 5.**
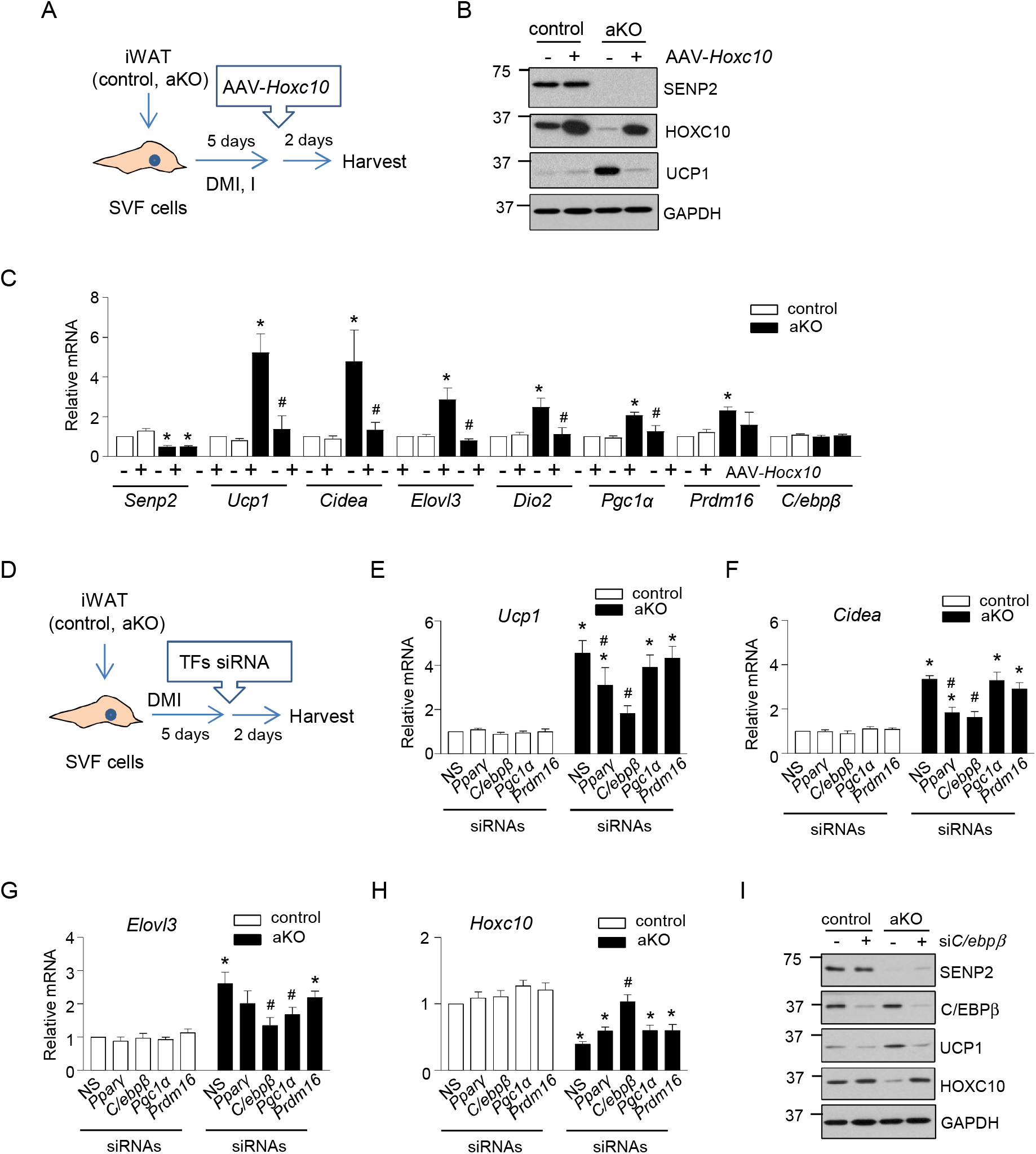
Suppression of HOXC10 stimulates differentiation of SENP2-deficient adipocytes toward beige adipocytes. (A-C) SVF isolated from iWAT of control or *Senp2-* aKO mice were differentiated to adipocytes for 5 days, and then infected with AAV-*Hoxc10* for 2 days. Cells were harvested and western blotting (B) and qPCR (C) were performed. The values of cells from control mice without AAV-*Hoxc10* treatment were set to 1, and the others were expressed as its relative values. \**P* < 0.05 vs control without AAV-*Hoxc10* by Student’s t-test, ^#^*P* < 0.05 vs aKO without AAV-*Hoxc10* by 2-way Anova, n = 4. (D-I) Cells were treated with siRNAs targeting the indicated transcription factors or siNS (non-specific) at day 5 of differentiation. Relative mRNA levels of *Ucp1* (E), *Cidea* (F), *Elovl3* (G) and *Hoxc10* (H) were determined. The values of cells from control mice treated siNS were set to 1 and the others were expressed as its relative values. \**P* < 0.05 vs control/siNS by Student’s t-test, ^#^*P* < 0.05 vs aKO/siNS by 2-way Anova, n = 4-5. (I) Western blotting was performed after si*C/ebp*β treatment. All data are represented as the mean ± SEM.

Next, we knocked down expression of the transcription factors PPARγ, C/EBPβ, PGC1α, and PRDM16 using specific siRNAs at day 5 of differentiation to investigate their involvement in SENP2–depletion–induced beige adipocyte differentiation (Fig. 5D and Supplemental Fig. 5B-5C). Although *Pparγ* knockdown affected expression of some brown adipocyte-specific genes such as *Ucp1*, *Cidea* and *Pgc1*α (Fig. 5E-5F and Supplemental Fig. 5C), clearer changes were observed with *C/ebp*β knockdown, which completely abrogated increased mRNA levels of *Ucp1, Cidea, Elovl3*, *Pgc1*α, and *Prdm16* and reduced *Hoxc10* mRNA induced by SENP2 depletion. In contrast, the effects of *Pgc1*α or *Prdm16* knockdown were not significant (Fig. 5E–5H and Supplemental Fig. 5C). The impact of *C/ebpβ* knockdown on UCP1 and HOXC10 was confirmed by western blot analysis (Fig. 5I), further highlighting C/EBPβ as a key factor involved in beige adipocyte differentiation induced by SENP2 deficiency in iWAT.

### Sumoylated C/EBPβ negatively regulates Hoxc10 transcription

Our results showed that suppression of *Hoxc10* was necessary for beige cell differentiation induced by SENP2 deficiency in iWAT and that C/EBPβ mediated *Hoxc10* suppression. Therefore, we next investigated how *Hoxc10* transcription is regulated by C/EBPβ. Several studies have reported that C/EBPβ can be sumoylated (35), which decreases its activity (36). C/EBPβ enhances promoter activities of various target genes; however, a few reports indicate that C/EBPβ also represses transcription of some target genes (37, 38). To test whether C/EBPβ directly suppresses *Hoxc10* transcription, we used transient transfection and a reporter assay system. Briefly, Cos7 cells were transfected with *Hoxc10* (−1567)-Luc and expression vectors for wild-type C/EBPβ (C/EBPβ WT) or C/EBPβ K133R, the latter of which contains a lysine-to-arginine mutation at the SUMO conjugation site, K133. Similar levels of C/EBPβ WT and C/EBPβ K133R were expressed in a dose-dependent manner (Supplemental Fig. 6). C/EBPβ WT, but not K133R, reduced the luciferase activity of *Hoxc10* (−1567)-Luc, and co-expression of SUMO and UBC9, a SUMO-conjugation enzyme, amplified the inhibitory activity of C/EBPβ WT (Fig. 6A). When SENP2 expression vectors were co-transfected, the C/EBPβ-mediated suppression of luciferase activity was diminished in a dose-dependent manner, whereas we observed no effect in C/EBPβ K133R (Fig. 6B). Sumoylated C/EBPβ was detected in cells transfected with C/EBPβ WT, but not K133R, and SUMO-conjugated C/EBPβ gradually decreased as SENP2 increased (Fig. 6C).

**Figure 6.**
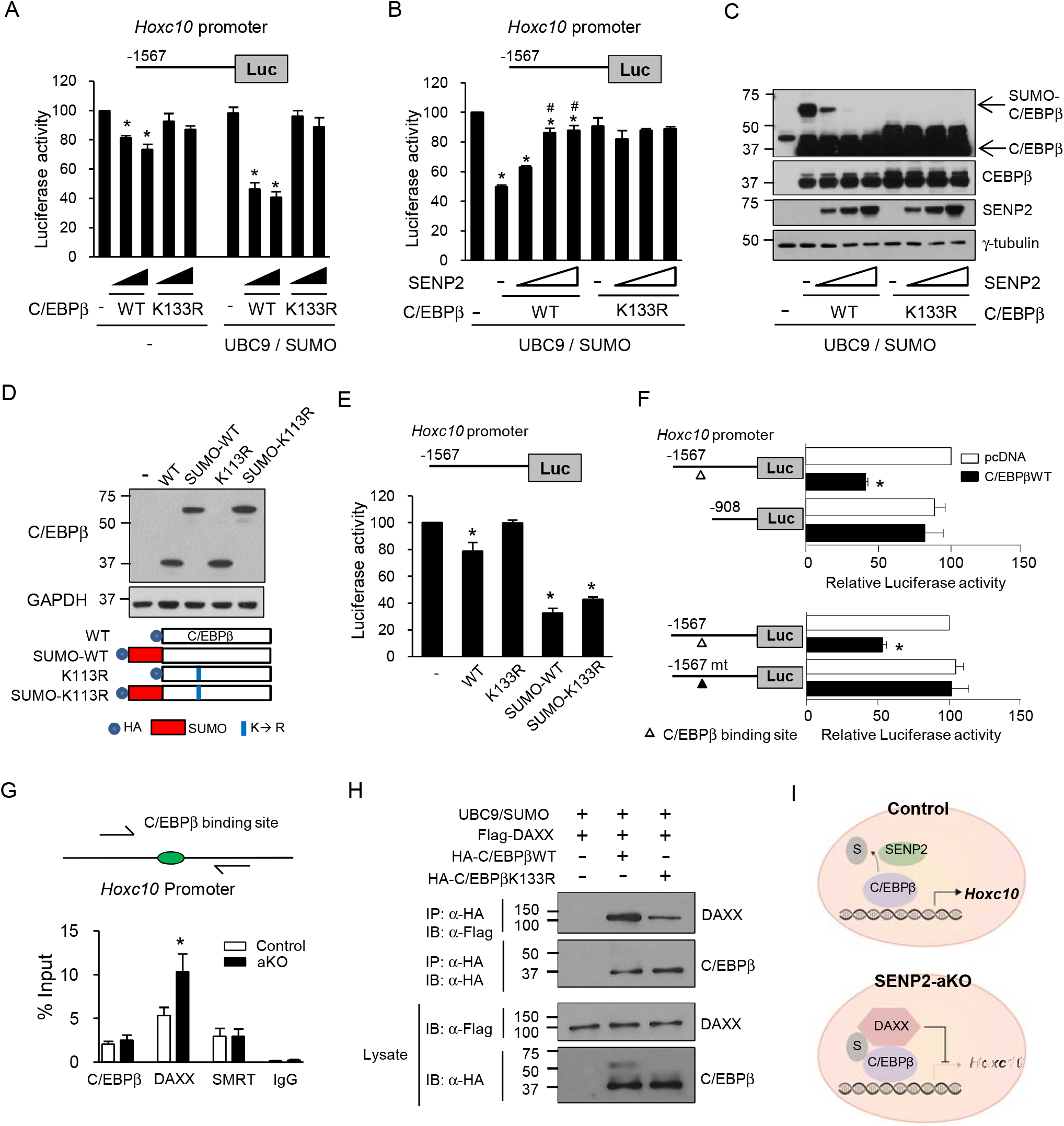
Inhibition of *Hoxc10* transcription by sumoylated C/EBPβ. (A) Cos 7 cells were transfected with *Hoxc10* (−1567 bp)-Luc, expression vectors for C/EBPβ WT or K133R, UBC9, SUMO and β-galactosidase. Luciferase activity of the cells transfected with *Hoxc10* (−1567 bp)-Luc without C/EBPβ was set to 100 and the others were expressed as its relative values. \**P* < 0.05 vs without C/EBPβ by Student’s t-test, n = 4. (B-C) Cells were transfected with different amounts of SENP2 expression vector. Luciferase activity (B) was measured. \**P* < 0.05 vs without C/EBPβ, ^#^*P* < 0.05 vs C/EBPβ WT without SENP2 by 2-way Anova. n = 3. Western blotting (C) was performed. Longer exposure of the anti-C/EBPβ blot shows sumoylated form of C/EBPβ (Top panel). (D-E) After transfection of C/EBPβ expression vectors and *Hoxc10*-Luc, western blot analysis (D) and luciferase assays (E) were performed. \**P* < 0.05 vs without C/EBPβ by Student’s t-test, n = 4. (F) Cells were transfected with different *Hoxc10*-Luc constructs. \**P* < 0.05 vs without C/EBPβWT by Student’s t-test, n = 5 (upper panel), n =3 (lower panel). (G) Adipocytes differentiated from iWAT SVF of control or *Senp2*-aKO mice were subjected for ChIP using an antibody of C/EBPβ, DAXX or SMRT. \**P* < 0.05 vs cells from control mice by Student’s t-test, n = 5. (H) Cos7 cells were transfected with the indicated expression vectors, and then immunoprecipitation was performed with an anti-HA antibody. All data are represented as the mean ± SEM. (I) A model showing the regulation of *Hoxc10* transcription by SENP2. All data are represented as the mean ± SEM.

To generate constitutive sumoylated C/EBPβ, we constructed expression vectors encoding SUMO fused to the N-terminus of C/EBPβ WT or K133R and confirmed expression of the SUMO-fused proteins by western blot analysis (Fig. 6D). SUMO-fused C/EBPβ K133R, as well as SUMO-fused C/EBPβ WT, effectively suppressed the *Hoxc10* promoter activity in the absence of UBC9 or SUMO overexpression (Fig. 6E). These results suggest that sumoylated C/EBPβ suppresses transcription of *Hoxc10*.

### Sumoylated C/EBPβ inhibits Hoxc10 transcription by recruiting the co-repressor death domain-associated protein (DAXX)

To identify a *cis*-acting element for C/EBPβ binding in the *Hoxc10* promoter, we generated a deletion construct of the *Hoxc10* promoter [*Hoxc*10 (−908)-Luc]. C/EBPβ WT did not suppress the luciferase activity of *Hoxc10* (−908)-Luc (Fig. 6F, upper panel). We induced mutations at a putative C/EBPβ binding site at −1030 [*Hoxc*10 (−1567mt)-Luc]. C/EBPβ WT did not inhibit the promoter activity of *Hoxc10* (−1567mt)-Luc (Fig. 6F, lower panel), demonstrating that C/EBPβ suppresses *Hoxc10* transcription through the C/EBPβ binding site.

C/EBPβ represses promoter activity of target genes through interaction with corepressors, such as silencing mediator of retinoid and thyroid hormone receptors (SMRT) (38, 39). Therefore, we hypothesized that sumoylated C/EBPβ recruits a corepressor to the transcriptional complex to inhibit transcription of *Hoxc10*. To test this hypothesis, we performed chromatin immunoprecipitation (ChIP) using adipocytes derived from iWAT SVF of control or *Senp2*-aKO mice with an antibody against SMRT. We also used an anti-DAXX antibody for ChIP, as DAXX has a SUMO-interacting motif and mediates SUMO-dependent transcriptional repression of glucocorticoid receptor and cAMP response element binding protein (40–42). Binding of Daxx, but not SMRT, to the C/EBPβ binding site of the *Hoxc10* promoter was stronger in SENP2-deficient adipocytes than in control adipocytes, without impacting C/EBPβ binding to the promoter (Fig. 6G). These results suggest that sumoylated C/EBPβ, which is increased by SENP2 depletion, recruits DAXX to suppress *Hoxc10* transcription. To confirm that sumoylated C/EBPβ efficiently interacts with DAXX, we performed co-immunoprecipitation and found that C/EBPβ WT interacted with DAXX to a greater extent than C/EBPβ K133R (Fig. 6H), suggesting that sumoylation of C/EBPβ promotes interaction between C/EBPβ and DAXX. In summary, SENP2 facilitates desumoylation of C/EBPβ, which results in high levels of *Hoxc10* transcription in control white adipocytes. However, when SENP2 is depleted, the sumoylated form of C/EBPβ increases and recruits DAXX, resulting in repression of *Hoxc10* transcription in iWAT adipocytes (Fig. 6I).

### SENP2 plays an important role in maintaining white adipocyte identity during adipogenesis through modulation of C/EBPβ in both iWAT and eWAT

We next confirmed the effect of SENP2 depletion on the differentiation of adipose progenitor cells using *Senp2* siRNAs (Fig. 7A). When SVF from iWAT of control mice was treated with siRNAs targeting *Senp2* at day 3 of differentiation, the mRNA levels of brown adipocyte-specific genes such as *Ucp1*, *Cidea*, and *Elovl3*, were significantly increased (Fig. 7B). We also detected a reduction of *Hoxc10* mRNAs. In addition, *Hoxc10* overexpression by infection of AAV-*Hoxc10* ameliorated the effects of *Senp2* knockdown (Fig. 7C and Supplemental Fig. 7A). We next tested whether SENP2 depletion also stimulates mature white adipocytes to acquire the features of beige adipocytes. When fully differentiated adipocytes (day 7 of differentiation) were transfected with siRNA targeting *Senp2*, we observed little or no change in the expression of brown adipocyte markers (Fig. 7B). In addition to iWAT browning, thermogenesis-related gene expression was elevated in eWATs of *Senp2*-aKO mice (Supplemental Fig. 3B and 3C), although the expression levels were lower than those in iWATs. We therefore tested whether *Senp2* knockdown induces expression of thermogenesis-related genes during differentiation of eWAT SVF (Fig. 7D). Similar to iWAT, *Senp2* knockdown by transfection with siRNA at day 3, but not at day 7, slightly increased *Ucp1* and *Cidea* expression (Fig. 7E). These results suggest that SENP2–depletion-induced beige or thermogenic adipocyte formation mainly occurs during differentiation of adipose progenitor cells rather than in mature white adipocytes.

**Figure 7.**
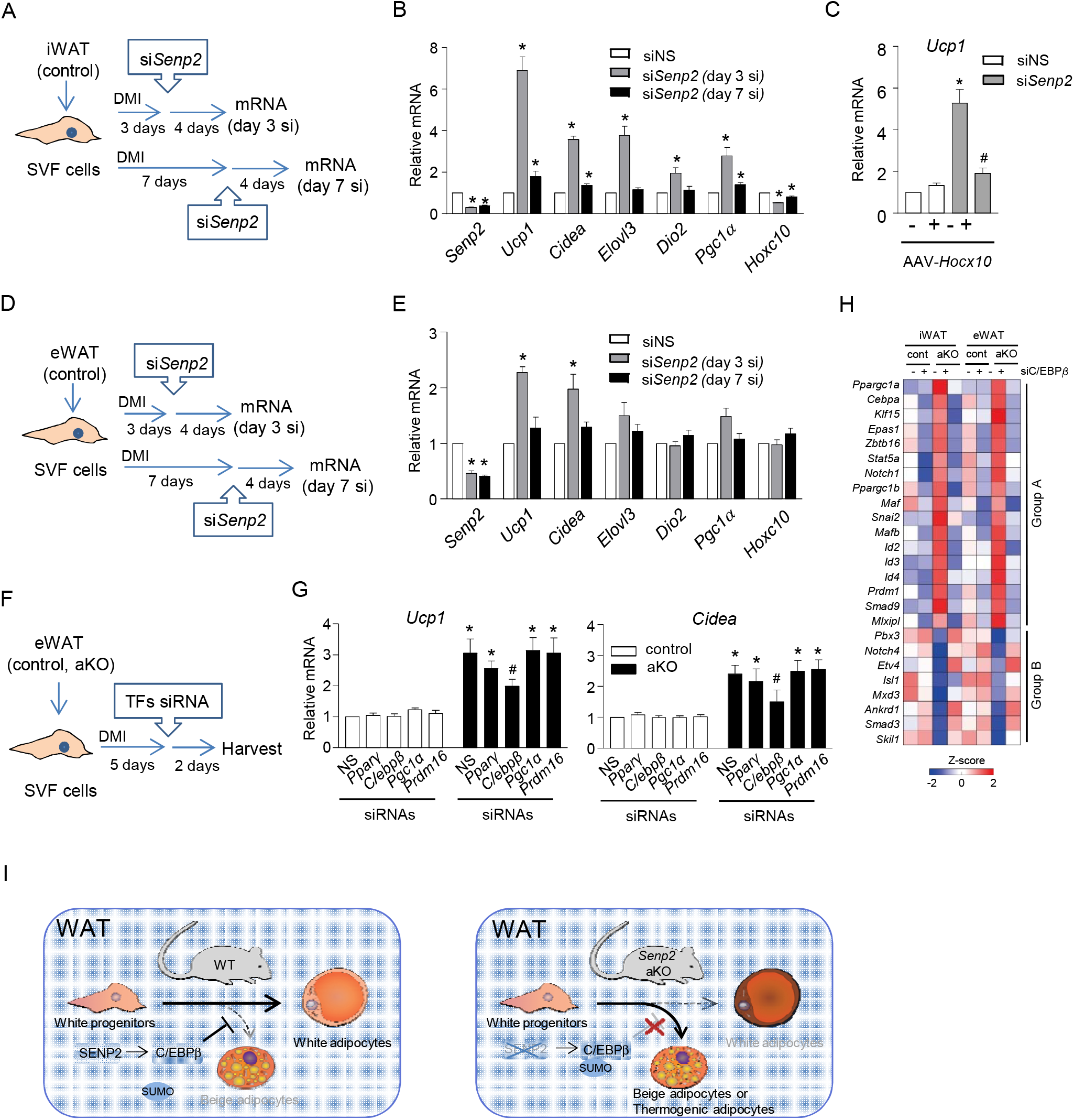
SENP2 depletion-induced thermogenic adipocyte differentiation through C/EBPβ in both iWAT and eWAT. (A-B) SVF isolated from iWAT of control mice was differentiated to adipocytes, and treated with siRNAs at day 3 or day 7 of differentiation. (B) qPCR was performed four days after siRNA treatment. The values of siNS-treated cells were set to 1. \**P* < 0.05 vs siNS by Student’s t-test, n = 5. (C) SVF cells, treated with siNS or si*Senp2* at day 3, were infected with AAV-*Hoxc10* at day 5 of differentiation. Two days after virus infection, cells were harvested and qPCR was performed. The values of siNS-treated cells without AAV-*Hoxc10* were set to 1. \**P* < 0.05 vs siNS without AAV-*Hoxc10*, ^#^*P* < 0.05 vs siSENP2 without AAV-*Hoxc10* by 2-way Anova, n = 5. (D-E) SVF was isolated from eWAT of control mice and treated with siRNAs. , n = 4. (F-G) eWAT SVF from control and Senp2-aKO mice was treated with the indicated siRNAs. eWAT SVF of control and *Senp2*-aKO mice were treated with siRNAs. The values of control mice/siNS treated cells were set to 1. \**P* < 0.05 vs siNS/control, ^#^*P* < 0.05 vs siNS/aKO by 2-way Anova, n = 4. All data are represented as the mean ± SEM. (H) Heat maps show the mean log_2_-fold changes of mRNA expression levels in the indicated conditions. (I) A model showing a function of SENP2 during differentiation of white adipose progenitor cells.

We also tested whether C/EBPβ mediates thermogenic gene induction by *Senp2* knockout in eWAT (Fig. 7F and Supplemental Fig. 7B). *Ucp1* and *Cidea* transcripts were increased in the adipocytes derived from eWAT SVF of *Senp2*-aKO mice, and knockdown of C/EBPβ suppressed those increases (Fig. 7G). These results demonstrate that C/EBPβ is also involved in thermogenic gene induction in eWAT of *Senp2*-aKO mice. Unlike iWAT, *Hoxc10* mRNA levels were very low in eWAT (Supplemental Fig. 3D), and *Hoxc10* transcription was not affected by *Senp2* knockout in eWAT (Supplemental Fig. 7B). Therefore, another factor that highly expressed in eWAT may act as a functional substitute for HOXC10.

To profile changes in the expression of various gene induced by SENP2 depletion in relation to C/EBPβ, we performed RNA sequencing after adipocyte differentiation of SVF from eWAT and iWAT of control or *Senp2*-aKO mice in the presence or absence of C/EBPβ knockdown (Supplemental Fig. 8A). We focused on expression patterns of various transcription factors. Several transcription factors whose expression was increased in both *Senp2*-aKO eWAT and iWAT in a C/EBPβ-dependent manner, such as PGC1α, STAT5, ZBTB16, and NOTCH1 (Fig. 7H, group A), have been reported to positively regulate beige adipocyte formation and thermogenesis (43–45). In addition to *Pgc1*α, transcription of *Stat5* and *Zbtb16* was also suppressed by *Hoxc10* overexpression in iWAT adipocytes (Supplemental Fig. 8B), which supports the C/EBPβ-HOXC10-transcription factor axis for mediating browning in iWAT. In contrast, expression of some transcription factors, such as SMAD3, NOTCH 4, and SKIL, was decreased in eWAT and iWAT of *Senp2*-aKO but was recovered by knockdown of C/EBPβ (Fig. 7H, group B). Interestingly, these transcription factors have been reported to promote white adipocyte differentiation and inhibit thermogenesis (46, 47). Therefore, we suggest that C/EBPβ is a master regulator of the expression of many transcription factors involved in adipocyte differentiation and browning in both iWAT and eWAT and that the direction of differentiation between white and beige or beige-like adipocytes can be determined by modulating SUMO modification of C/EBPβ. In other words, SENP2 plays a role in maintaining white adipocyte identity during differentiation of white adipose progenitor cells by modulation of C/EBPβ (Fig. 7I).

## Discussion

We have previously shown that SENP2 plays an important role in the early stages of adipocyte differentiation *in vitro*. To identify other possible functions of SENP2 in adipocytes, we generated adipocyte-specific *Senp2* knockout mice using an adiponectin-*Cre* system in the current study. Here, we demonstrated that SENP2 depletion significantly reduced fat accumulation and facilitated formation of beige adipocytes in WAT. In addition, SENP2 depletion during adipogenesis caused white adipose progenitor cells to differentiate toward beige adipocytes. Therefore, SENP2 has dual functions in white adipocytes: regulating initial adipogenesis and maintaining white adipocyte identity throughout differentiation. Interestingly, we identified C/EBPβ as a direct target of SENP2 that modulates both of these functions. SENP2 stabilized C/EBPβ through desumoylation at the early stages of adipogenesis (33) and abolished sumoylated C/EBPβ-mediated suppression of browning inhibitors, such as *Hoxc10*, which maintained white adipocyte identity during differentiation.

Traditionally, C/EBPβ induces transcription of various target genes, although some reports have described C/EBPβ-mediated transcriptional repression. Several co-repressors, including SMRT, SWItch/Sucrose Non-Fementable, and CA150, interact with C/EBPβ and repress target gene expression (38, 39, 48). In this study, we demonstrated that a co-repressor, DAXX, interacted with C/EBPβ and repressed *Hoxc10* transcription. DAXX has a known SUMO-interacting motif (42), and we observed that the DAXX binding of C/EBPβ was increased by C/EBPβ sumoylation. Therefore, we suggest that sumoylated C/EBPβ effectively recruits DAXX, resulting in suppression of *Hoxc10* transcription (Figure 6I). Further experiments will be required to determine whether the C/EBPβ-DAXX complex represses transcription of other genes whose expression was downregulated in white adipocytes of *Senp2*-aKO mice.

In addition to *Hoxc10*, expression of several other transcription factors was modulated in WAT by *Senp2* knockout. Although our data demonstrated that *Hoxc10* suppression was important for browning in iWAT of *Senp2*-aKO mice, other transcription factors may also contribute to browning. As such, the transcription of several *Hox* genes, including *Hoxc6*, *Hoxc8*, and *Hoxb5*, was also downregulated in iWAT of *Senp2*-aKO mice (Supplemental Fig. 3A). Notably, these genes are highly expressed in white adipocytes, and that HOXC8 is known to inhibit brown adipocyte differentiation (49). Therefore, several HOX factors may work together to regulate expression of genes that determine brown or white adipocyte differentiation. Although the mechanism by which HOXC10 inhibits beige adipocyte differentiation in WAT has not been studied in detail, our RNA-seq analysis (Fig. 7H and Supplemental Fig. 8B) showed that HOXC10 inhibits expression of several transcription factors involved in beige adipogenesis, such as *Pgc1*α and *Stat5a,* suggesting that HOXC10 suppresses expression of upstream transcription factors, rather than directly regulating expression of thermogenic genes, such as *Ucp1*. However, further promoter analyses are required to determine how HOXC10 regulates expression of these transcription factors. Unlike iWAT, *Hoxc10* expression is very low in eWAT; therefore, a different C/EBPβ downstream factor(s) may suppress thermogenic activation in eWAT.

Our results suggest that the direction of adipocyte differentiation can be controlled through post-translational modification of C/EBPβ in iWAT and eWAT even though iWAT and eWAT adipocytes originate from different cell lineages. WAT consists of adipocytes derived from several types of precursor cells, while beige adipocyte differentiation is induced in specific lineages (20, 50, 51). Therefore, determining which cell lineage(s) is induced to differentiate into beige adipocytes as a result of *Senp2* knockout is of great interest. C/EBPβ is known to be involved in both white and brown adipocyte differentiation (22) and is thought to regulate the expression of *Ucp1* and other thermogenic genes by directly binding to their promoters (52). Therefore, these varied and sometimes contradictory roles of C/EBPβ may be accomplished by precise modulation of C/EBPβ through post-translational modification such as sumoylation.

One study demonstrated that SENP2 deficiency reduced adipose lipid storage and increased fat accumulation in other tissues, such as the liver and muscle tissues, which resulted in insulin resistance (53). With the exception of decreased fat mass, however, those results were different from our data due to deficiency of browning effects. In that study, *Senp2*-aKO mice were generated by deleting exons 13 and 14 from *Senp2*, which encode the C-terminus of SENP2. In our construct, however, exon 3 was deleted, and we confirmed that no truncated form of SENP2 was produced in adipose tissue of *Senp2*-aKO mice (Supplemental Fig. 1C). Therefore, the contradictory results of the two studies could be a consequence of different knockout strategies.

BAT mass was dramatically reduced in *Senp2*-aKO mice fed either a normal chow diet or HFD. We observed enlarged lipid droplets in BAT of HFD-fed Senp2-aKO mice, suggesting that the function of brown fat in *Senp2*-aKO mice was impaired. A recent report demonstrated that SENP2 is important for brown adipose development using *Senp2*-BAT knockout mice generated with myogenic factor 5-*Cre* (54). Therefore, SENP2 depletion might directly affect brown adipocyte differentiation and function, as *Senp2* transcription was downregulated in BAT of *Senp2*-aKO mice. However, BAT reduction in *Senp2*-aKO mice may also be a secondary effect of beige adipocyte accumulation in WAT. Further study using brown adipocyte-specific *Senp2* knockout mice generated using *Ucp1*-*Cre* will clarify the function of SENP2 in mature brown adipocytes.

To summarize, SENP2, highly expressed in white adipocytes, inhibits browning of WAT by modulation of C/EBPβ. Inducing browning of WAT by inactivation of SENP2 or downregulation of *Senp2* expression can be a therapeutic approach to ameliorate obesity and metabolic syndromes including diabetes.

## Materials and Methods

### Generation of adipocyte-specific Senp2 knockout mice

*Senp2^flox/+^* mice were generated by the inGenious Targeting Laboratory (Stony Brook, NY, USA) as described previously (55). *Senp2* adipocyte specific-knockout (*Senp2*-aKO) mice were generated by a serial mating *Senp2^flox/flox^* with adiponectin-*Cre* transgenic mice (Jackson lab).

### Mice and Metabolic analysis

All aspects of animal care and experiments were conducted in accordance with the Guide for the Care and Use of Laboratory Animals of the National Institutes of Health, and approved by the Institutional Animal Care and Use Committee of Seoul National University Bundang Hospital, Korea (Permit Number: BA1403-149/013-01). All animals were housed at 22-24°C, with a 12:12-h light-dark cycle and ad libitum access to standard pelleted chow or high-fat diet, (HFD; 58 kcal% fat w/sucrose, D12331; Research Diets, New Brunswick, NJ, USA) and water. Eight week-old mice were fed a HFD for 12 weeks. Body weight was measured weekly, and body composition was measured by dual-energy X-ray absorptiometry (DXA) and LUNAR Prodigy scanner with software version 8.10 (GE Healthcare, PA, USA). Mice were maintained at room temperature or 4°C when indicated, and body temperatures were measured rectally using a digital thermometer. Metabolic cage experiments were conducted using a Comprehensive Lab Animal Monitoring System (CLAMS; Columbus Instruments, Columbus, OH, USA). Food and water intake were recorded weekly during HFD feeding. For the glucose tolerance test (GTT), glucose (1 g/kg body weight) was treated by intraperitoneal (i.p.) injection after 16 h starvation. For the insulin tolerance test (ITT), mice fasted for 6 h were injected with human insulin (1 U/kg body weight). Serum glucose levels were determined using an OneTouch Ultra glucometer (LifeScan, Milpitas, CA, USA). Serum insulin (ALPCO, NA, USA), leptin, (Millipore, Billerica, MA, USA) and adiponectin (Millipore) were measured by enzyme-linked immunosorbent assays. Triacylglycerol (Cayman Chemical, Ann Arbor, MI, USA) and free fatty acid (Wako chemicals, Richmond, VA, USA) levels were determined using colorimetric assays.

### Histological analysis

Tissues were fixed in 4 % formaldehyde, embedded in paraffin and sectioned. Sections were subjected to hematoxylin and eosin (H&E) staining. Immunohistochemistry was performed with antibodies against UCP1 (Abcam, Cambridge, UK). Lipid droplet diameters were measured using Leica software. 100-150 adipocytes per mouse were measured.

### Microarray experiments

The mRNA profiles of iWATs from control and *Senp2*-aKO mice were analyzed using the Agilent *mouse* 8×60k chip, which includes 62,976 probes corresponding to 21,895 annotated genes (Agilent, Santa Clara, CA, USA). Gene expression profiles were generated from four replicates in each condition. Total RNA was extracted, and the integrity of the total RNA was analyzed using an Agilent 2100 Bioanalyzer. RNA integrity numbers for all samples were larger than 7.3. The *in vitro* transcription was then performed to generate cRNA, which was hybridized onto each array, according to the manufacturer’s instructions. The array was scanned using a SureScan Microarray Scanner (Agilent).

### Identification of differentially expressed genes (DEGs)

The intensities of the probes from the arrays were converted to log_2_-intensities and then normalized using the quantile normalization method (56). To identify DEGs, an integrative statistical hypothesis tests was performed as previously described (57). Briefly, for each gene, a T-statistic value was calculated using Student’s t-test and also a log_2_-fold change in the comparison of *Senp2*-aKO versus control was calculated. We then estimated empirical distributions of T-statistic values and log_2_-fold changes for the null hypothesis (i.e. the genes are not differentially expressed) by performing all possible random permutations of the eight samples. Using the estimated empirical distributions, we computed adjusted p-values for the two tests for each gene and then combined these p-values with Stouffer’s method (58). Finally, DEGs was identified as the genes with combined p-values ≤ 0.01 and absolute log_2_-fold changes ≥ 0.58 (1.5-fold change).

### Functional enrichment analysis

Functional enrichment analysis was performed using DAVID software to identify gene ontology biological processes (GOBPs) enriched by the DEGs (59). The GOBPs enriched by the up- and down-regulated genes were identified as the ones with the enrichment P-value < 0.05 calculated by DAVID software.

### mRNA sequencing and data analysis

Total RNA was obtained from WATs (eWAT and iWAT) of control or *Senp2*-aKO mice in the absence or presence of *C/ebp*β knockdown. The integrity of the total RNA was analyzed using an Agilent 2100 Bioanalyzer. RNA integrity numbers for all samples were larger than 8.9. Poly(A) mRNA isolation from total RNA and subsequent fragmentation were performed using the NEBNext Ultra II Directional RNA-seq Kit, according to the manufacturer’s instructions (New England Biolabs, Ipswich, MA, USA). The adaptor ligated libraries were sequenced using an Illumina HiSeq X Ten (Ebiogen, Korea). mRNA sequencing was performed for three independent replicates under each condition. From the resulting read sequences for each sample, adapter sequences (TruSeq universal and indexed adapters) were removed using the cutadapt software (version 2.7). The remaining reads were then aligned to the *Mus musculus* reference genome (GRCm38) using TopHat2 software (version 2.1.1) with default parameters (60).After the alignment, we counted the numbers of reads mapped to the gene features (GTF file of GRCm38.91) using HTSeq (61).Read counts for the samples in each condition were then normalized using the trimmed mean of M-values (TMM) normalization method in the edgeR package (62).

### Identification of the differential expression patterns by non-negative matrix factorization (NMF) clustering

To effectively identify the major clusters of genes showing altered expression patterns in multiple comparisons, we performed NMF clustering as previously described (63). For NMF analysis, we calculated log_2_-fold changes of each gene in the following comparisons using the normalized read counts in eWAT or iWAT samples: si*C/ebp*β cont/siNS_cont, siNS_*Senp2*-aKO/siNS_cont, and si*C/ebp*β_*Senp2*-aKO/siNS_cont. These log_2_-fold changes were combined into a fold change matrix. Using the fold change matrix, we then performed NMF clustering and selected the genes defining the individual clusters with the following parameters as previously described (63): the number of clusters (*k*) = 50 and p-value cutoff = 0.01 for the gene selection.

### Real time qPCR

Total RNAs were isolated using TRIzol (Invitrogen, MA, USA) according to the manufacturer’s instruction. Real-time qPCR was performed (in duplicates) using the SYBR-master mix (Takara, Otsu, Shiga, Japan) and the ABI 7500 Real-time PCR system (Applied Biosystem, CA, USA). Each cycle threshold (Ct) value was subtracted from Ct value of GAPDH or β-actin (△Ct), and then subtracted from the value of each control set (^△△^Ct). Relative mRNA levels were expressed as 2^−ΔΔCt^. Primer sequences are listed in Supplemental Table 1.

### Isolation of SVF and in vitro differentiation

Stromal vascular fraction (SVF) cells were isolated from the iWAT of 8 to12 week-old mice. Tissues were minced in digestion buffer (100 mmol/l HEPES pH7.4, 120 mmol/l NaCl, 50 mmol/l KCl, 5 mmol/l glucose, 1 mmol/l CaCl_2_, 1.5 % BSA-fatty acid free, and 1 mg/ml collagenase II), and incubated at 37°C with shaking for 1 h. Large particles were removed using a 100 μm cell strainer, and 20 ml Hank’s buffered salt solution (HBSS, Gibco Life Technolosies, Carlsbad, CA, USA) supplemented with 10 % fetal bovine serum (FBS, Gibco Life Technolosies) and 10 mM HEPES was added to filtrates. The filtrates were centrifuged at 2,500 rpm for 10 min to remove floating adipocytes. The pelleted SVF cells were suspended in Dulbecco’s modified Eagle’s medium (DMEM, Gibco Life Technolosies) supplemented with 10 % FBS, and seeded in 12-well plates. Adipocyte differentiation of confluent SVF cells was induced by treatment with DMI (1 μmol/l dexamethasone, 0.5 mmol/l isobutylmethyxanthine and 0.1 mg/ml insulin) in DMEM supplemented with 10 % FBS. Two days after the treatment, cells were maintained in DMEM supplemented with 10 % FBS and 1 μg/ml insulin for additional 5 days. The media was changed every 2 days.

### Oil Red O staining

Cells were washed once with PBS, and subsequently fixed for 10 min in 4 % formalin. After fixation, lipid droplets were stained by 5 % Oil Red O in 60 % isopropanol for 1 h. Cells were washed twice with distilled water and observed using a microscope.

### Transfection of siRNAs, and infection of AVV

Small interfering RNAs (siRNAs) of *Senp2*, *C/ebpβ*, *Pparγ*, *Pgc1α*, and *Prdm16* were purchased from Dharmacon (Chicago, IL, USA). Nonspecific siRNAs (siNS) were purchased from BIONEER (Daejeon, Korea). The siRNAs (100 nmol/l) were mixed with RNAiMAX (Invitrogen) in 100 μL of serum-free DMEM and incubated for 30 min at room temperature. The complex was treated to cells with 400 μL of serum-free DMEM. After 3 h incubation, 500 μl of DMEM supplemented with 10 % FBS was added. Adeno-associated virus (AAV)-Flag-*Hoxc10* was previously described (32). AAV-Flag-*Hoxc10* was purified by AAV Mini Purification Kit (BioVision, Milpitas, CA, USA). Purified virus was diluted in Tris-HCl buffer at 1/100 ratio and then treated to cells.

### Plasmids and antibodies

A DNA fragment from −1567 bp to −22 bp (from ATG) of the *Hoxc10* promoter was inserted into the pGL2-basic vector (Promega, Madison, WI) to generate *Hoxc10* (−1567)-Luc. In addition, *Hoxc10* (−908)-Luc was constructed by inserting the promoter fragment from −908 bp to −22 bp of the mouse *Hoxc10* promoter into pGL2-enhancer vector. To generate *Hoxc10* (−1567 mt)-Luc, DNA sequences between −1034 bp and −1026 bp of the *Hoxc10* (−1567)-Luc, 5′-TGGAGCAAAG-3′, were changed to 5′-ACCTGCAAAG −3′ by using the Quick-Change site directed mutagenesis Kit (Stratagene, La Jolla, CA, USA). Expression vectors for SENP2, C/EBPβ WT, C/EBPβ K133R, SUMO and UBC9 were described previously (33). The expression vector for FLAG-DAXX was a gift from Dr. C.H. Chung (Seoul National University). To generate SUMO-tagged C/EBPβ, cDNA of SUMO1 was ligated between HA and C/EBPβ in pcDNA-HA-C/EBPβWT or K133R. Antibodies against SENP2, HOXC10, DAXX, SMRT and GAPDH were purchased from Santa Cruz Biotechnology (Santa Cruz, CA, USA). Antibodies against C/EBPβ and UCP1 were obtained from Abcam (Cambridge, UK), and the γ-tubulin antibody and CL316243 were purchased from Sigma-Aldrich (St Louis, MO, USA).

### Transient transfection and luciferase reporter assay

COS7 cells were cultured in DMEM supplemented with 10 % FBS. COS7 cells were seeded at 12-well plates and transfected with reporter vectors (0.3 μg), expression vector for C/EBPβ WT, C/EBPβ K133R (0.01 μg or 0.025 μg), UBC9 (0.1 μg), SUMO (0.1 μg), SENP2 (0.01, 0.05 and 0.1 μg) and pRSV-β-galactosidase (0.1 μg) using Lipofectamine and Plus reagent (Invitrogen). Luciferase and β-galactosidase activities were measured according to the manufacturer’s instruction (Promega, Madison, WI).

### Immunoprecipitation and Western blot analysis

For immunoprecipitation, COS7 cells plated on 6-well plates were transfected with expression vectors for HA-C/EBPβ WT or HA-C/EBPβ K133R (0.25 μg), Flag-DAXX (1 μg), Myc-SUMO (0.5 μg) and Flag-UBC9 (0.5 μg) by using Lipofectamin plus (Invitrogen). After incubation for 24 h, cells were lysed in the RIPA buffer (Millipore) supplemented with protease inhibitors (10 μg/μl aprotinin, 10 μg/μl leupeptin and 1 mM PMSF). Cell lysates (0.5 mg) were incubated with anti-HA antibody-coupled agarose beads (Roche, Basel, Switzerland) for 16 h at 4 °C. The precipitates were washed five times and subjected to SDS– polyacrylamide gel electrophoresis. For western blot analysis of tissue or cell lysates, samples were prepared in RIPA buffer supplemented with protease inhibitors. Proteins on SDS– polyacrylamide gel were transferred onto a nitrocellulose membrane (GE Healthcare). The bands were visualized utilizing Enhanced Chemiluminescence (Pierce, Rockford, IL, USA).

### Chromatin IP linked qPCR

SVF was isolated from iWAT of control or *Senp2*-aKO mice, and then differentiated to adipocytes for 7 days. After cross-linking and DNA fragmentation, nuclear extracts were immunoprecipitated with anti-C/EBPβ, anti-DAXX, anti-SMRT or control IgG antibodies. Real-time qPCR was performed using the primers (Supplemental Table 1) for the C/EBPβ binding site (−1030 bp) of the *Hoxc10* promoter.

### Statistical analysis

Statistical analysis was performed using SPSS version 12.0 (SPSS inc.). Statistical significances of the differences were determined by Student’s *t* test and 2-way ANOVA. *P* < 0.05 was considered statistically significant.

## Acknowledgements

This research was supported by Basic Science research Program through the National Research Foundation of Korea (NRF) funded by the Ministry of education (NRF-2015R1D1A1A09059551) (NRF-2016R1A2B3010373) (NRF-2019R1A2C3009517) and by the National Institutes of Health (R01DK111529 to YBK).

## Author contributions

K.S.P. and S.S.C. designed the study, interpreted the data and wrote the manuscript. J.S.L., J.N., Y.D.K., and S.L. performed the experiments and analyzed the results. S.C. and D.H. performed transcriptomic analysis. Y.B.K., W.H. and Y.J.P. interpreted the results and edited the manuscript.

## Declaration of interests

The authors declare no competing interests.

## Supplemental Figures

**Supplemental Figure 1. Generation of adipocyte-specific** *Senp2* **knockout mice** (A) SENP2 protein levels in several tissues, gastronecmius (GM), liver and WATs, were compared by western blot analysis. (B) *Senp2* mRNA levels in adipose tissues were measured by qPCR. The values of control mice were set to1. \**P* < 0.05 vs control by Student’s t-test, n = 13-14. (C) After fractionation of adipocytes and stromal vascular fraction (SVF) from WAT, the lysates were subjected to western blotting of SENP2. (D) Representative western blots showing SENP2 protein levels in gastronecmius muscle (GM) and liver.

**Supplemental Figure 2. Tissue mass and serum parameters of control and** *Senp2***-aKO mice on a HFD** (A) Average food intake during HFD feeding. n = 5–7. (B) After 14 weeks of HFD feeding, bone mass density (BMD) and bone mass were measured. n = 9-11. (C) Weight of liver and GM tissues. n = 9-11. (D) Serum insulin, (E) serum triacylglycerol (TG) and free fatty acids (FFA) after HFD feeding. n = 16-22. (F) Serum leptin and adiponectin levels. \**P* < 0.05 by Student’s t-test, n= 21-22. All data are presented as mean ± SEM.

**Supplemental Figure 3. Gene expression profiles in iWAT and eWAT of control and** *Senp2***-aKO mice**. (A) Heat map showing the mRNA levels of transcription factors in iWAT of HFD-fed mice. n = 4. (B-C) Relative mRNA levels of the indicated genes in eWAT of mice on a NCD (B) or HFD (C). The values obtained from control mice were set to 1. \**P* < 0.05 by Student’s t-test, n = 13-15. (D) Relative mRNAs of *Senp2, C/ebpb* and *Hoxc10* in adipose tissues n = 3. All data are presented as mean ± SEM.

**Supplemental Figure 4. Expression of thermogenic genes during differentiation of SENP2-defiecent cells** Relative mRNA levels of *Cidea*, *Elovl3* and *Pgc1a*. The values of control cells at day 0 were set to 1 and the others were its relative values. \**P* < 0.05 vs control by 2-way Anova, n = 3. All data are represented as the mean ± SEM.

**Supplemental Figure 5. Knockdown of several transcription factors during differentiation of adipose progenitor cells** (A) *Hoxc10* mRNA levels were determined 2 days after AAV-*Hoxc10* infection. The values of the control cells not treated with AAV-*Hoxc10* were set to1. \**P* < 0.05 vs control without AAV-*Hoxc10* by Student’s t-test, ^#^*P* < 0.05 vs aKO without AAV-*Hoxc10* by 2-way Anova, n = 4. (B) Knockdown of *Pparγ* and *C/ebp*β was confirmed. (C) The mRNA levels of *Senp2*, *Pgc1a* and *Prdm16* were analyzed after knockdown of the transcription factors. The value of the control cells treated with siNS was set to1. \**P* < 0.05 vs control/siNS by Student’s t-test, ^#^*P* < 0.05 vs aKO/siNS by 2-way Anova, n = 4-5. All data are represented as the mean ± SEM.

**Supplemental Figure 6. Expression of C/EBP**β **after transfection** Cos7 cells were transfected with C/EBPβ expression vectors (5, 10 and 25 ng), pcDNA-HA-C/EBPβWT and pcDNA-HA-C/EBPβ K133R. Expression of C/EBPβ was confirmed by western blot analysis.

**Supplemental Figure 7. Effects of overexpression of** *Hoxc10* **on beige adipogenesis of SENP2-deficient progenitor cells** (A) iWAT SVF cells, treated with siNS or si*Senp2* at day 3, were infected with AAV-*Hoxc10* at day 5 of differentiation. The values of siNS-treated cells without AAV-*Hoxc10* were set to 1. \**P* < 0.05 vs siNS without AAV-*Hoxc10*, ^#^*P* < 0.05 vs siSENP2 without AAV-*Hoxc10* by 2-way Anova, n = 5. (B) eWAT SVF were treated with siRNAs. The values of control /siNS treated cells were set to 1. \**P* < 0.05 vs siNS/control, ^#^*P* < 0.05 vs siNS/aKO by 2-way Anova, n = 4. All data are represented as the mean ± SEM.

**Supplemental Figure 8. Profiles of genes showing C/EBPβ-dependent expression changes in both eWAT and iWAT of** *Senp2***-aKO** (A) Heat maps with respect to the mean expression levels in control (Cont_siNS), for genes in the clusters showing C/EBPβ-mediated up-regulation (A; Clusters 6, 16, 20, 30) and down-regulation (B; Clusters 19, 21, 22, 25) in eWAT and iWAT. Cellular processes enriched by the up- (Group A) and down-regulated (Group B) genes are also shown in the bar plots and the enrichment significance (p-value) was displayed as -log_10_(p). (B) *Hoxc10* was overexpressed by infection of AAV-*Hoxc10* in iWAT SVF-derived adipocytes, and qPCR was performed. The values of cells from control mice without AAV-*Hoxc10* treatment were set to 1, and the others were expressed as its relative values. \**P* < 0.05 vs control without AAV-*Hoxc10* by Student’s t-test, ^#^*P* < 0.05 vs aKO without AAV-*Hoxc10* by 2-way Anova, n = 4.

## Notes

### Competing Interest Statement

The authors have declared no competing interest.

### Summary of Updates

Figure 1 K-->J typing error corrected; Acknowledgements updated; Figure files (PPT) uploaded

