## Supplementary figures and images for "SUMO-specific protease 2 (SENP2) suppresses browning of white adipose tissue through C/EBPβ modulation"

### supple Fig 1.jpeg

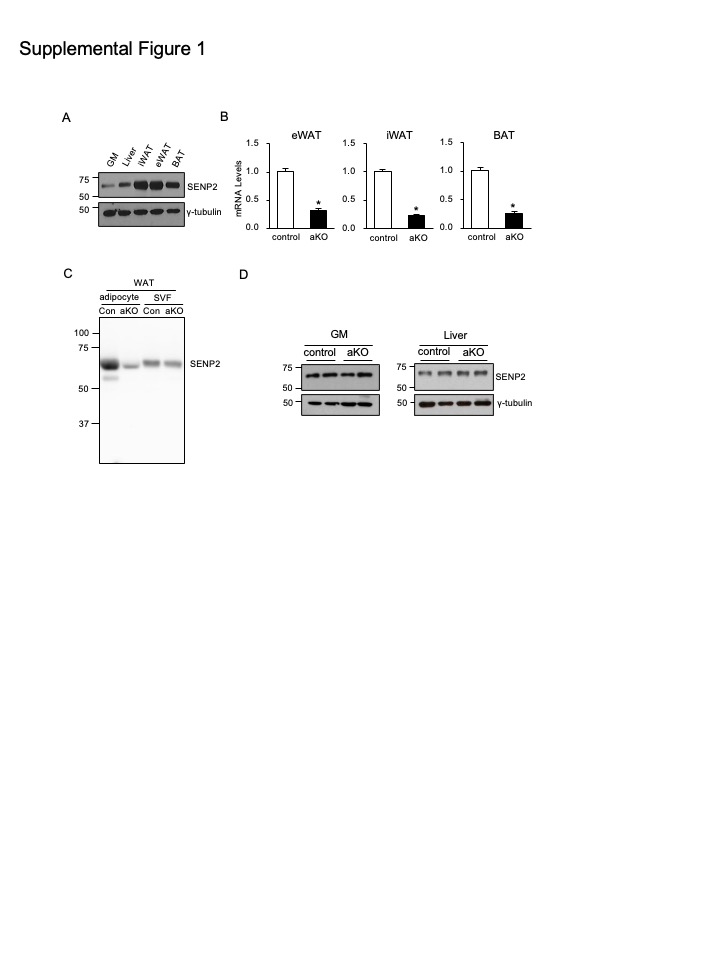

### supple Fig 2.jpeg

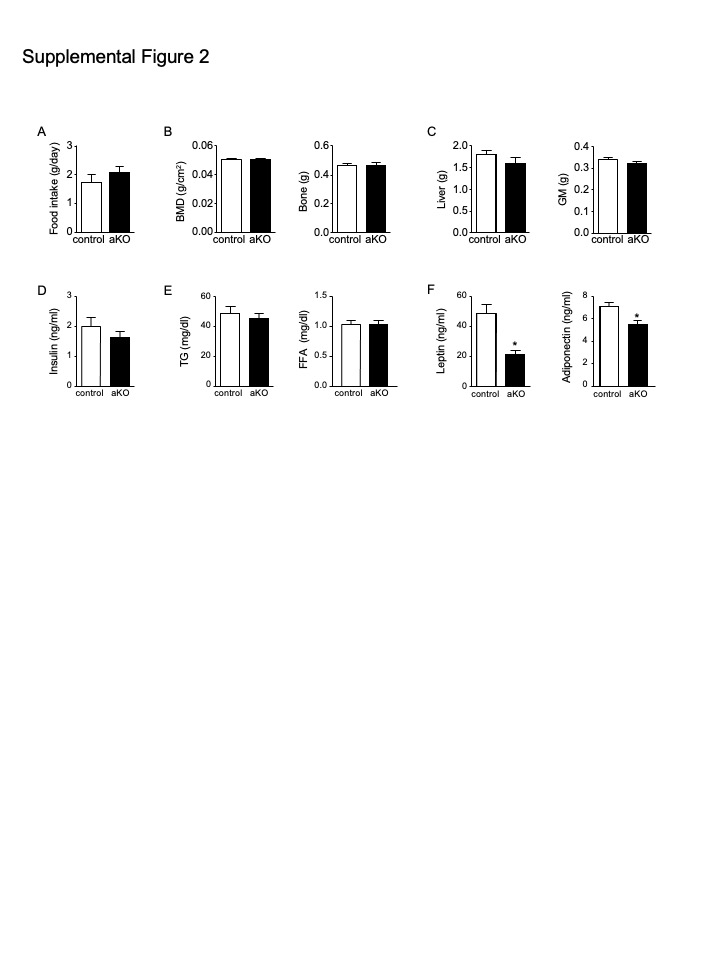

### supple Fig 3.jpeg

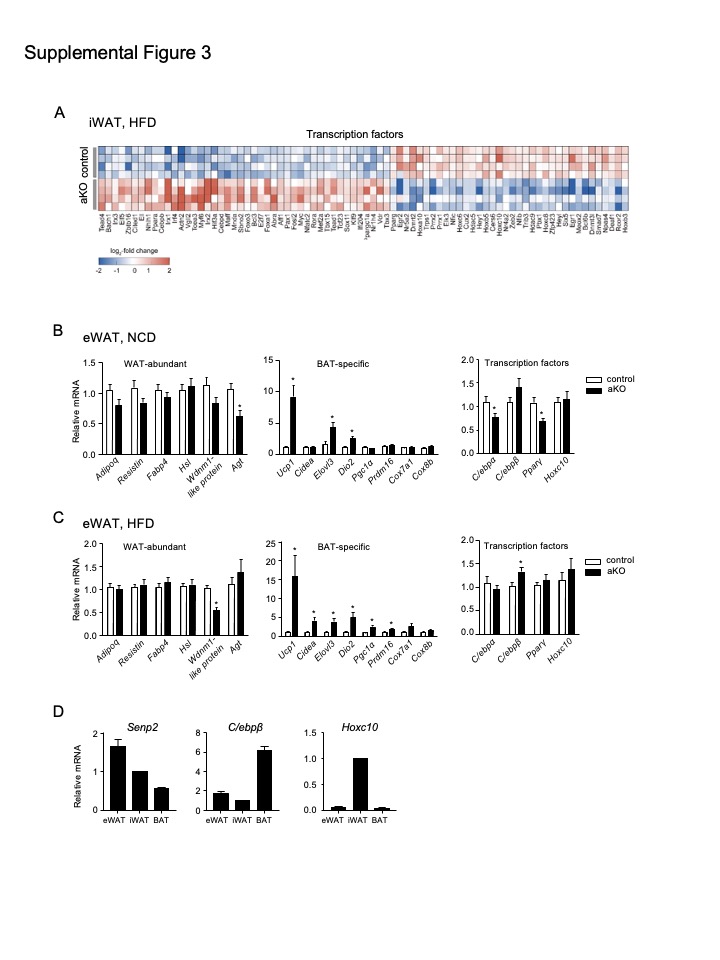

### supple Fig 4.jpeg

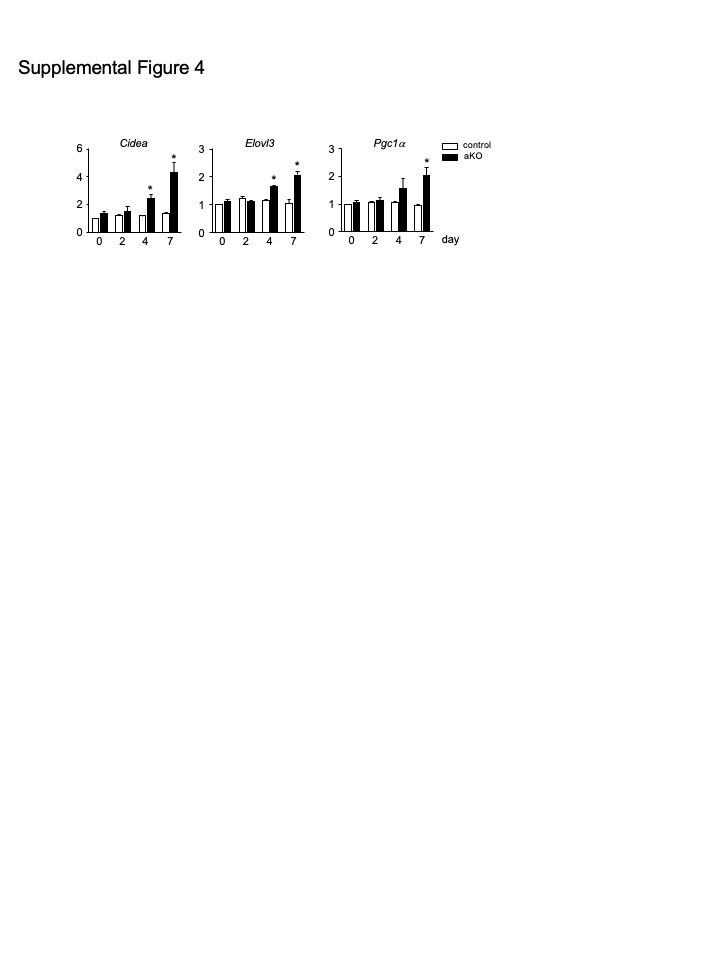

### supple Fig 5.jpeg

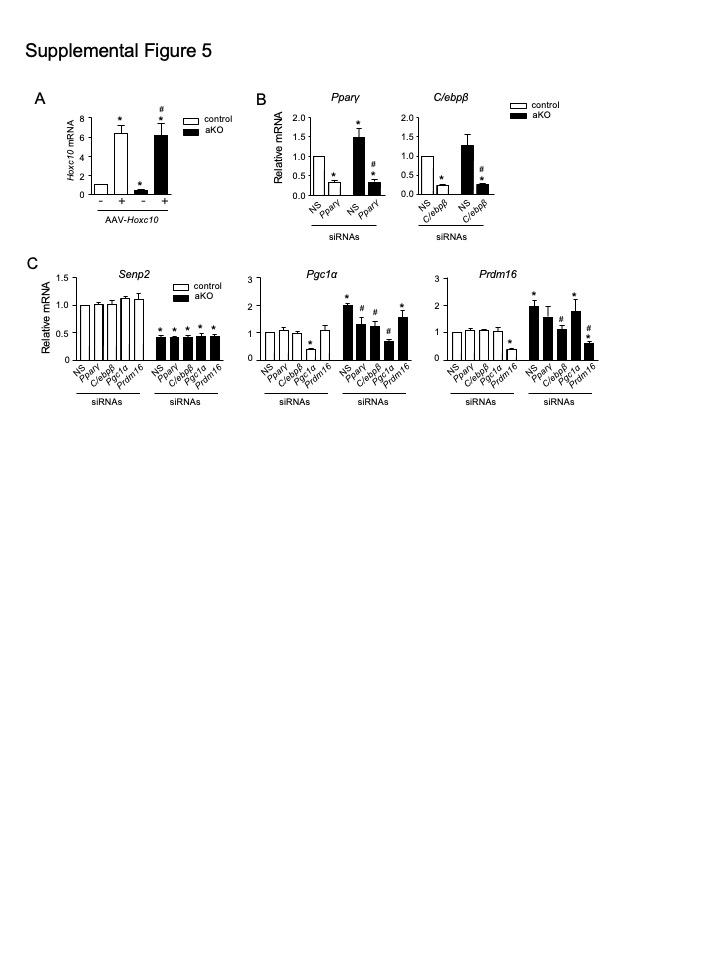

### supple Fig 6.jpeg

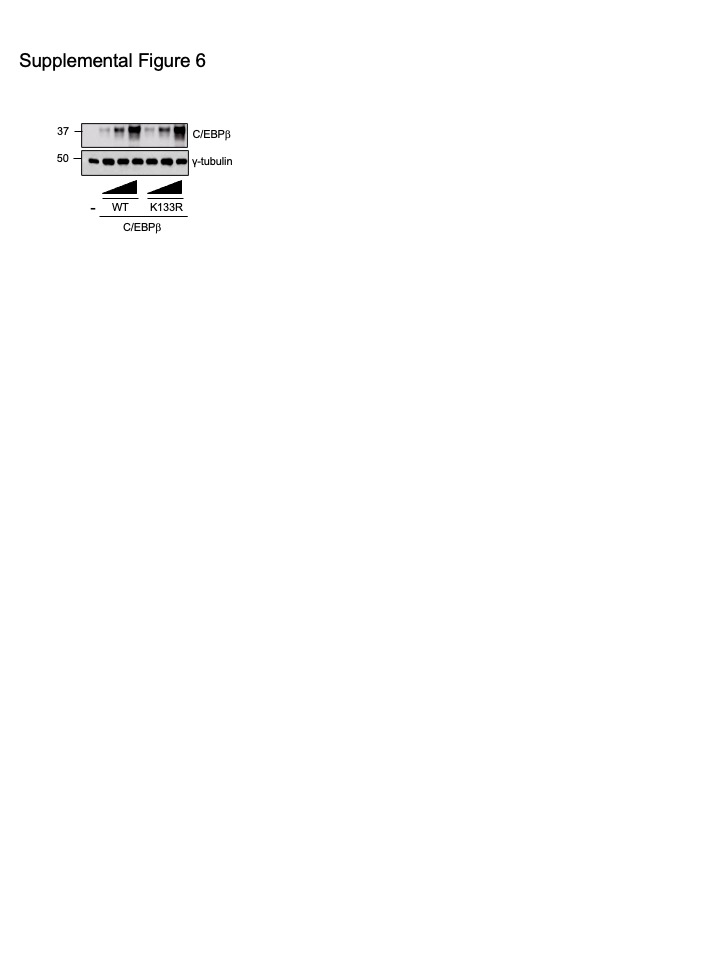

### supple Fig 7.jpeg

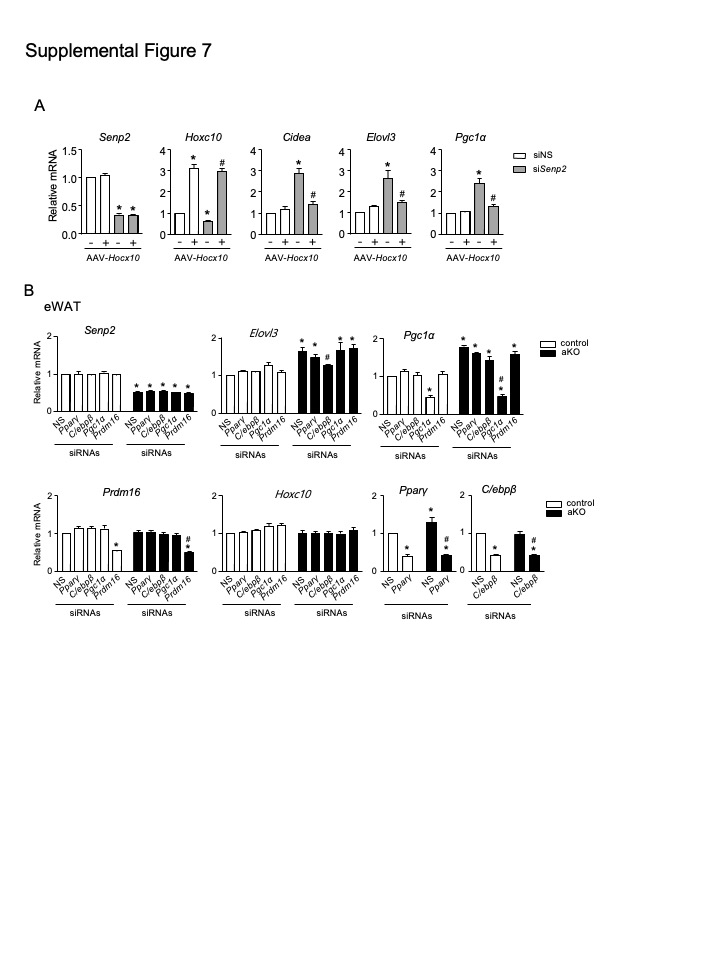

### supple Fig 8.jpeg

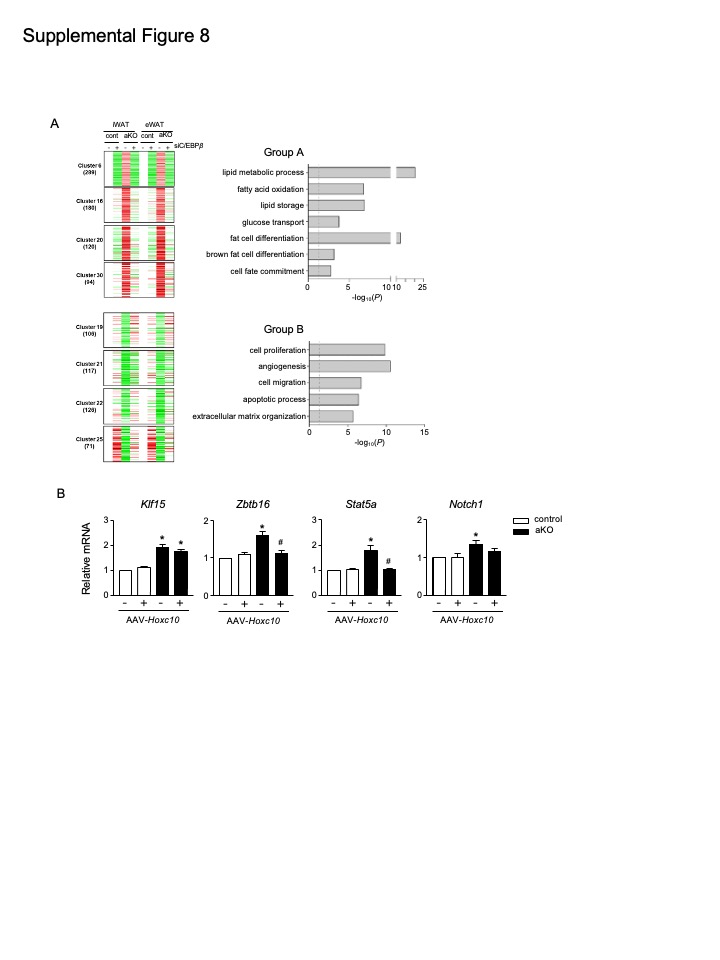

### supple Table 1.jpeg

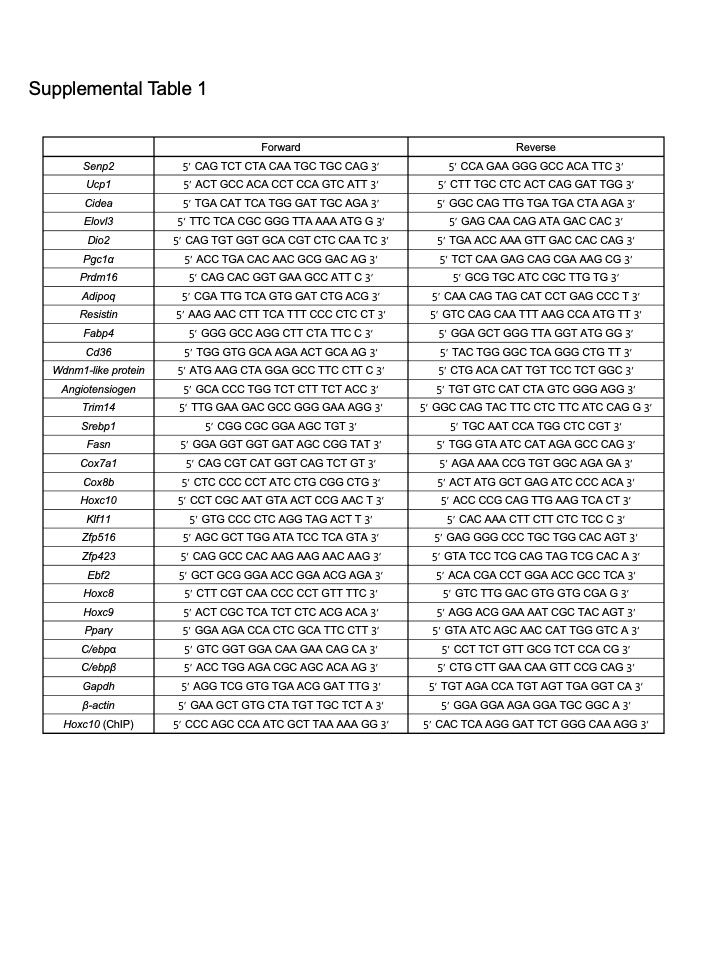
